# Ageas enables time-agnostic cell fate inference from single-cell and spatial multi-omics data

**DOI:** 10.64898/2026.08.30.748098

**Authors:** Junyao Jiang, Alex Kong, Gang Yu

**Affiliations:** School of Life Sciences, Westlake University, Hangzhou, China; Danish Research Institute of Translational Neuroscience (DANDRITE), Nordic EMBL Partnership for Molecular Medicine, Aarhus University, Aarhus, Denmark; Department of Biomedicine, Aarhus University, Aarhus, Denmark; Institute of Modern Biology, Nanjing University, Nanjing, China; Gene Center and Department of Biochemistry, Ludwig-Maximilians-Universität München, Munich, Germany

**Author notes:** These authors contributed equally to this work (Junyao Jiang, Gang Yu).

## Abstract

Understanding cell fate decisions is fundamental to developmental biology and disease research. However, experimental lineage tracing requires genetic manipulation, which is impractical in many systems, particularly in humans. Computational approaches often rely on time-resolved measurements, which single-cell and spatial omics studies rarely provide. Here, we present Ageas, a time-agnostic transfer learning framework for cell fate inference from single-cell and spatial multi-omics data. Ageas seeks to address these limitations by learning fate memory from terminal cell populations and transferring this information to progenitor or intermediate cells, allowing fate bias inference from static molecular snapshots. To support generalization across molecular modalities, Ageas employs a data-adaptive ensemble strategy with automated model selection. On benchmark datasets with lineage-traced single-cell transcriptomic and epigenomic profiles, as well as spatial transcriptomics, Ageas achieves performance comparable to or exceeding existing methods. Applying Ageas to a 3D human embryo, we identify an anterior–posterior trend of epiblast fate priming, with anterior epiblast cells biased toward ectodermal fates and posterior cells toward primitive streak-derived lineages, accompanied by regionally graded fate-associated regulatory programs. Together, these results support Ageas as a general framework for inferring cell fate decisions from static molecular snapshots.

## 1 Introduction

Characterizing the developmental history and fate decisions of cells is crucial for understanding both normal development and disease progression. Recent advancements in single-cell and spatial omics technologies, including single-cell RNA sequencing (scRNA-seq), single-cell Assay for Transposase-Accessible Chromatin with sequencing (scATAC-seq), and spatial transcriptomics (ST), have enabled high-resolution profiling of cellular states across tissues[1–3]. A major limitation of these techniques is that they capture only static snapshots of cellular and molecular states, lacking the mapping information for cell fates from progenitors to differentiated progeny[4].

One approach to address the limitations of snapshot data has been the development of prospective genetic lineage tracing technologies[4, 5]. These methods integrate heritable DNA barcoding with single-cell or spatial sequencing, so that progenitor cells are uniquely labeled and their clonal descendants can be identified at later time points[6–11]. Such lineage-tracing strategies have yielded valuable insights into lineage segregation events in model organisms by linking early and late cell states. However, current prospective genetic lineage tracing requires introducing synthetic sequences into the genome, posing serious challenges for many systems, particularly in human tissues (Fig. 1A)[4, 12].

**Figure 1:**
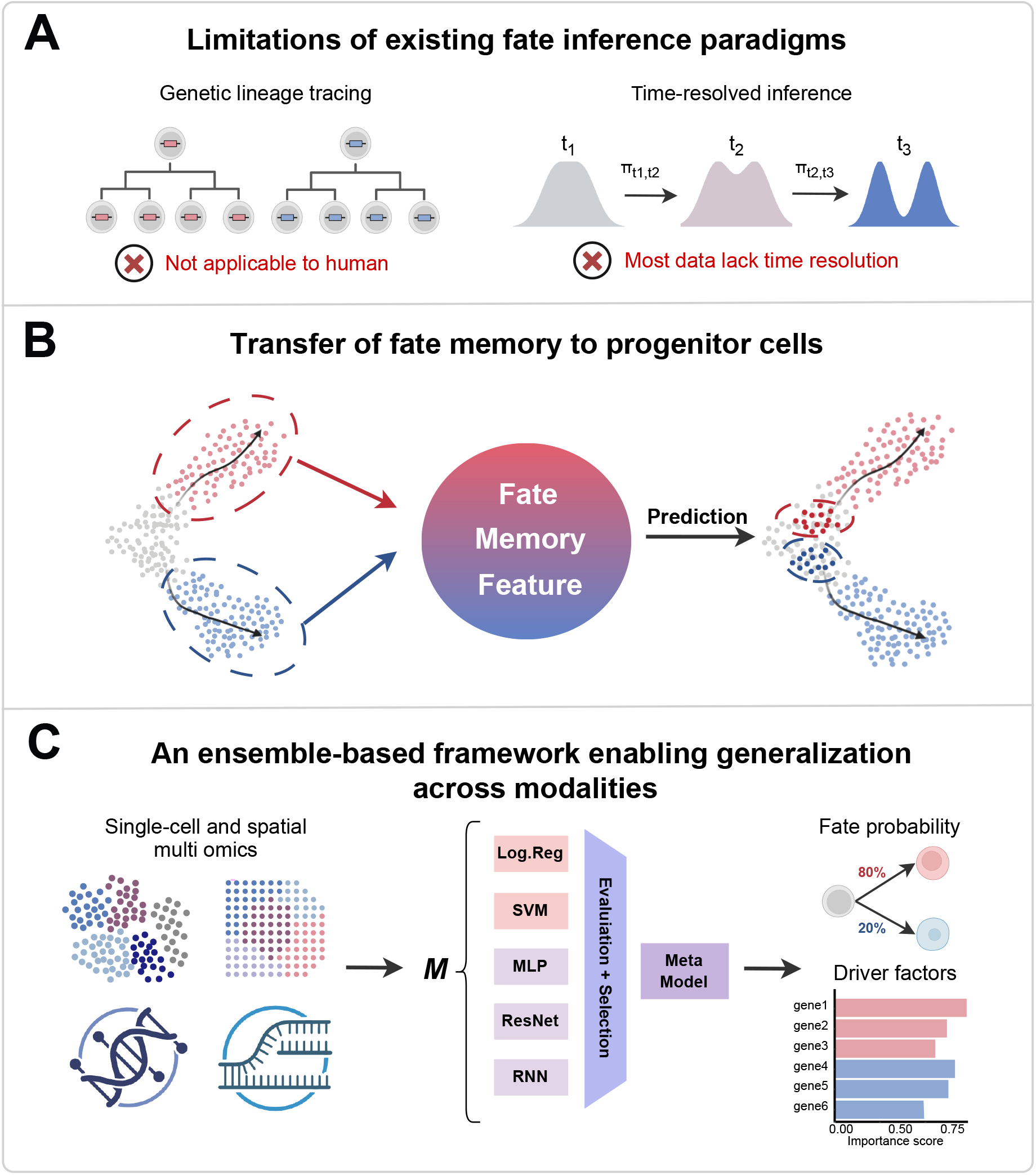
The Ageas framework for time-agnostic cell fate inference. **A,** Schematic contrasting the limitations of genetic lineage tracing and time-resolved computational inference. **B,** Ageas learns fate-discriminative memory from terminal states and transfers it to progenitors to predict fate from static snapshots. **C,** The ensemble-based architecture. Ageas aggregates multiple classifiers into a meta-model to ensure generalization across single-cell and spatial omics modalities.

Meanwhile, several computational methods have been developed to map cell fates and predict progenitor fate biases from gene expression data. Waddington-OT[13] (WOT) uses optimal transport (OT) to infer cell state transitions across time points while accounting for cell proliferation and death. CoSpar[14] infers cell dynamics by integrating coherent but sparse lineage signals to resolve early fate biases. CellRank[15, 16] models cell state transitions as a Markov chain and computes fate probabilities toward terminal states with various transition kernels, including RNA velocity and pseudotime. Despite their proven utility, these computational approaches are optimized with time-resolved experimental designs (e.g., multi-timepoint sampling). However, many biologically important studies are limited to single-timepoint snapshots. This includes large-scale studies of human development, such as human embryo and fetal atlases[17–19], as well as cross-sectional single-cell analyses of cancer progression, including oligodendroglioma, head and neck cancer, and glioblastoma[20–22].

Furthermore, many current computational frameworks model transcriptional dynamics, with less attention to the broader regulatory landscape underlying cell fate decisions. Lineage commitment could also be shaped by epigenetic mechanisms, such as chromatin accessibility[23, 24], which can prime regulatory potential and influence developmental trajectories before overt transcriptional divergence occurs[10, 25–27]. Together, these gaps motivate computational strategies that can infer cell fate biases across diverse molecular modalities.

Here, we introduce Ageas, a time-agnostic transfer learning framework that approaches cell fate inference as a problem of information transfer rather than temporal reconstruction. By extracting fate-discriminative molecular signatures from differentiated progeny and projecting them onto progenitor populations, Ageas infers fate bias from static cellular snapshots without requiring time-resolved measurements. To support generalization across diverse molecular modalities, Ageas employs a data-adaptive model selection and evaluation module that constructs an ensemble tailored to each dataset. This strategy aims to facilitate interpretable fate inference across single-cell and spatial multi-omics data. Applying Ageas to transcriptomic, epigenomic, and spatial datasets, we show that cellular fate biases can be inferred from isolated cross-sections, and we evaluate this capability on datasets with lineage-tracing ground truth as well as in spatial regeneration systems. Specifically, Ageas infers spatially organized fate priming within the human epiblast, identifying an anterior-posterior gradient of ectoderm and primitive streak potential, alongside fate-associated molecular programs that complement insights from experimental lineage tracing. Collectively, these findings suggest Ageas as a time-independent framework for inferring cell fate trajectories and their associated molecular drivers directly from static single-cell and spatial multi-omics data.

## 2 Results

### 2.1 Overview of Ageas

Ageas is a transfer learning framework designed to infer cell fate bias from single-cell or spatially resolved omics data, bypassing the need for time-course experiments or genetic lineage tracing. Instead of calculating transition maps or reconstructing developmental trajectories, Ageas reconceptualizes fate inference as the transfer of fate-specific molecular memory. Specifically, the framework extracts fate-discriminative molecular signatures from mature, differentiated cells and projects this information onto progenitor populations to forecast lineage outcomes (Fig. 1B). In practice, an ensemble of supervised classification models is trained on terminal cell states with defined identities, identifying critical fate-memory features, such as distinct transcriptional profiles and chromatin accessibility landscapes, that segregate divergent lineages. Subsequently, this ensemble is applied to progenitor or intermediate cells to directly predict the probability of specific lineage commitment based on static molecular snapshots, operating entirely independently of temporal cellular relationships.

To ensure broad applicability across diverse molecular modalities, Ageas implements a data-adaptive strategy leveraging automated machine learning (AutoML) principles[28, 29], systematically evaluating a wide search space of model configurations spanning various algorithmic architectures (Fig. 1C). Crucially, the framework decouples the identification of optimal hyperparameter configurations from the final instantiation of the predictive models. Candidate models are first initialized using the top-performing configurations and subsequently trained on the defined terminal cell states. Only fully fitted models that exceed a stringent empirical performance threshold are retained, ultimately aggregating into the final predictive ensemble based on their respective fate probability outputs. This architectural design not only ensures broad generalizability across heterogeneous datasets but also renders Ageas inherently modality-agnostic. By completely detaching fate prediction from any specific molecular readout, the framework operates seamlessly across single-cell transcriptomic, epigenomic, and spatial expression profiles (Fig. 1C). Beyond estimating fate probabilities, Ageas yields deep biological interpretability via an integrated explanation module that computes feature importance scores. This module highlights the genes, regulatory elements, or other features most associated with each lineage outcome, thereby identifying key lineage drivers and molecular programs linked to fate decisions from static data.

### 2.2 Assessment of Ageas classification performance as a foundation for fate inference

The Ageas framework utilizes an ensemble of multiple models to classify terminal cell types and capture lineage-specific, fate-memory signatures. To establish its predictive reliability, we systematically benchmarked Ageas against established classification algorithms across four representative scRNA-seq datasets[30–33] characterized by varying data sizes and cellular complexities (Supplementary Table 1). Specifically, we benchmarked Ageas against four widely adopted methods, representing both correlation-based heuristics (Seurat[34] and SingleR[35]) and deep learning-based architectures (scANVI[36] and TOSICA[37]).

To mitigate stochastic variance and ensure robust performance metrics, we implemented a rigorous five-fold cross-validation (5-CV) strategy for each dataset (Supplementary Fig. 1A). Across all benchmarked cohorts, Ageas consistently demonstrated classification state-of-the-art level performance. Notably, in the highly heterogeneous colon dataset[33], Ageas achieved a median accuracy of 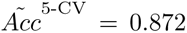 with scANVI achieving the second best performance of 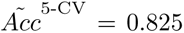 (Supplementary Fig. 1A,B). This improvement over the classification performance underscores the enhanced sensitivity of Ageas to capturing subtle yet fate-associated transcriptional features within complex and heterogeneous cell populations.

We further challenged the model using the Zheng68K peripheral blood mononuclear cell (PBMC) dataset[30], a rigorous benchmark characterized by pronounced class imbalance and high transcriptional similarity among closely related immune subtypes. Under these conditions, Ageas maintained robust classification performance with a median accuracy of 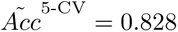 and a median macro F1 score of 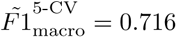, marginally exceeding that of scANVI 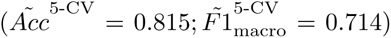 and outperforming traditional correlation-based approaches (Supplementary Fig. 1A,C).

Collectively, these benchmarking analyses demonstrate that Ageas provides stable and high-fidelity classification across diverse cellular landscapes, establishing a rigorous foundation for downstream developmental fate inference.

### 2.3 Benchmarking cell fate inference performance of Ageas

We evaluated the predictive fidelity of Ageas for cell fate inference by benchmarking its performance against three established algorithms[13–15] (CoSpar, Waddington-OT, and Cell-Rank) using two independent single-cell lineage-tracing datasets[6, 25].

The first benchmark leveraged an induced endoderm progenitor (iEP) reprogramming dataset[25] (Fig. 2A,B), which maps the direct lineage reprogramming of mouse embryonic fibroblasts (MEFs) and captures a developmental bifurcation into either a successfully reprogrammed iEP trajectory or a off-target, dead-end state. To evaluate model performances, we focused on models’ predictive ability to classify the terminal fate bias of day-3 (D3) progenitors relying exclusively on their static gene expression profiles. As the result, Ageas exhibited superior performance across all evaluation metrics, with accuracy of *Acc* = 0.873; macro F1 score of *F*1_macro_ = 0.870; and the area under the receiver operating characteristic curve (AUROC) of *AUROC* = 0.88 (Fig. 2C,D, Supplementary Fig. 2A, and Supplementary Table 2). This performance exceeded the next best method (CoSpar) which achieved *Acc* = 0.751; *F*1_macro_ = 0.767; and *AUROC* = 0.832. These performance gains across complementary metrics indicate that Ageas more precisely resolves fate-determining signals at the progenitor stage.

**Figure 2:**
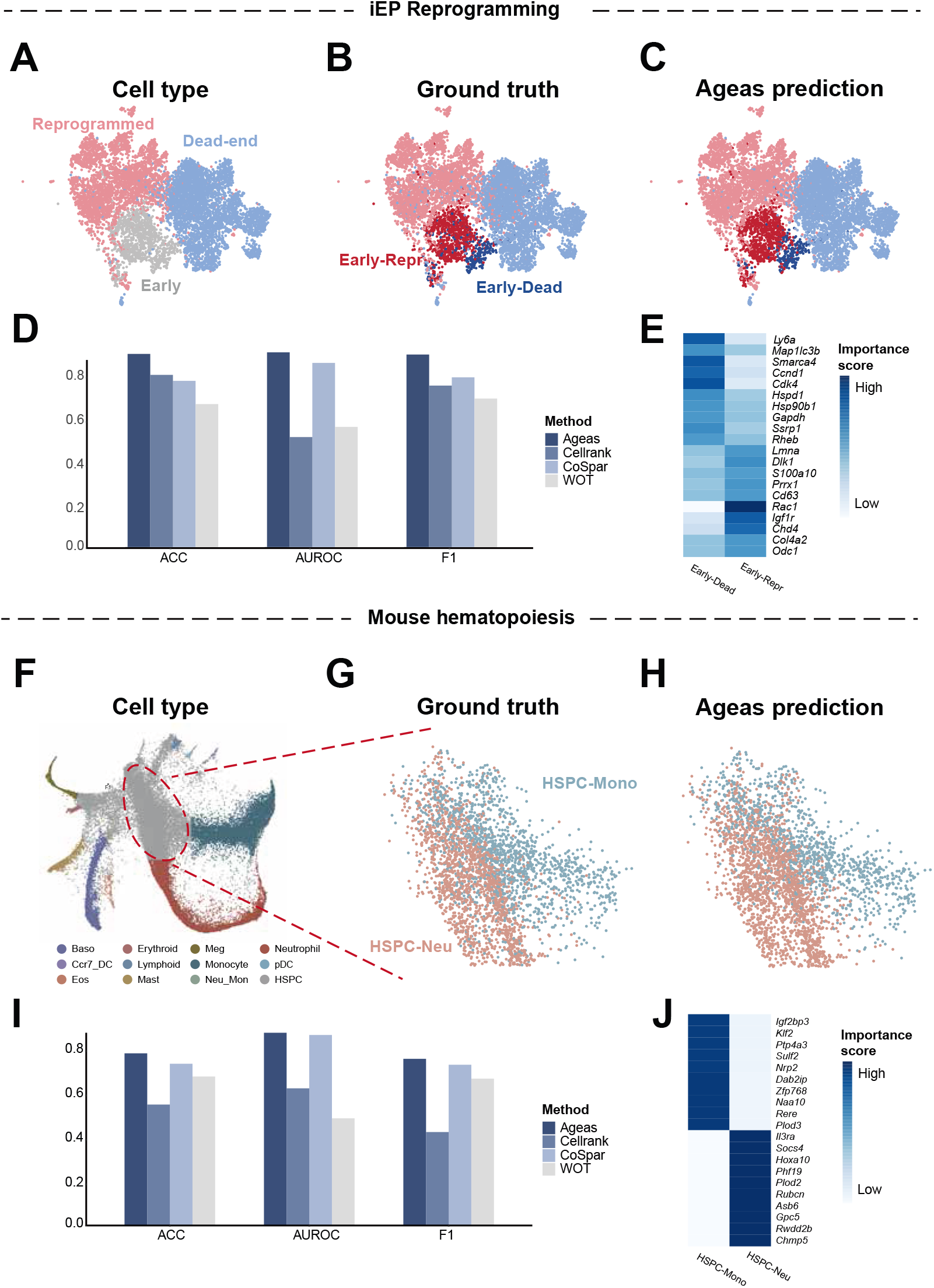
Benchmarking Ageas on single-cell lineage-tracing datasets. **A,** UMAP visualization of cell types in iEP reprogramming dataset. **B,** Ground truth obtained from lineage tracing experiments (CellTag-multi), distinguishing *Early-Reprogramming* (red) from *Early-Dead* (blue) cells. **C,** Ageas-predicted fate bias for early cells. **D,** Performance comparison against CellRank, CoSpar, and WOT. **E,** Heatmap showing the top lineage driver genes identified by the Ageas explanation module, associated with *Early-Dead* and *Early-Reprogramming* fates. **F,** SPRING visualization of hematopoietic progenitor differentiation. **G,** Ground truth for HSPCs toward Monocyte (HSPC-Mono) or Neutrophil (HSPC-Neu) fates. **H,** Ageas-predicted fate bias for HSPCs. **I,** Performance metrics comparing Ageas with other methods. **J,** Heatmap showing the top lineage driver genes identified by the Ageas explanation module, associated with HSPC-Mono and HSPC-Neu lineages.

Leveraging the explainability module integrated within the Ageas framework, we subsequently identified the primary transcriptomic drivers underpinning these cell fate predictions. Within the successfully reprogrammed progenitor cohort (*Early-Reprogramming*), Ageas identified the upregulation of *Ly6a*, *Cdk4*, and *Rheb*, which aligns with known programs of enhanced cellular plasticity[38, 39] and proliferation[40]. In contrast, the progenitor cohort fated for a dead-end state (*Early-Dead*) was characterized by elevated expression of *Lmna*, *Dlk1*, and *Prrx1*, signatures associated with mesenchymal identity[41, 42] and unsuccessful cellular reprogramming[43]. These results confirm that Ageas effectively captures regulatory programs driving early fate divergence (Fig. 2E).

We further assessed algorithmic generalizability by extending our analysis to a mouse hematopoiesis dataset[6], in which hematopoietic stem and progenitor cells (HSPCs) differentiate into monocytic and neutrophilic lineages (Fig. 2F-G). Here, HSPCs were designated as early progenitors, with the differentiated populations serving as terminal outcomes. Ageas consistently yielded the highest accuracy (*Acc* = 0.790) and macro F1 scores *F*1_macro_ = 0.765, alongside a modest improvement in AUROC (*AUROC* = 0.882) compared to CoSpar (*AUROC* = 0.872), affirming its predictive advantage across highly divergent biological systems (Fig. 2H,I, and Supplementary Fig. 2B).

Furthermore, model explainability successfully isolated distinct lineage-associated genetic programs governing hematopoiesis (Fig. 2J). Cells biased toward the HSPC-neutrophil trajectory were marked by the expressions of *Klf2* [44], *Nrp2* [45], and *Ptp4a3* [46], genes critical for neutrophil-specific differentiation. In contrast, cells exhibiting an HSPC-monocyte bias are identified by the expression of *Hoxa10* [47], *Phf19* [48], and *Plod2* [49], factors heavily implicated in progenitor maintenance and monocytic commitment.

Together, these findings demonstrate that Ageas reliably infers progenitor fate biases from static gene expression data, outperforming current state-of-the-art frameworks, while simultaneously identifying biologically coherent regulatory programs that dictate lineage commitment.

### 2.4 Ageas generalizes to epigenomic modalities and infers cell fate bias from static chromatin accessibility profiles

Cell fate decisions are orchestrated by both transcriptional states and underlying epigenomic configurations, which prime regulatory potential well before overt gene expression divergence occurs. Given the modality-agnostic architecture of the Ageas ensemble framework, we hypothesized that its predictive capacity would seamlessly extend to chromatin accessibility landscapes. To test this, we applied Ageas to a scATAC-seq dataset with ground truth from a CellTag-multi reprogramming experiment[25] incorporating lineage-traced ground truth from a CellTag-multi lineage tracing system (Fig. 3A,B). Operating on time-agnostic peak-by-cell accessibility matrices, Ageas successfully predicted early fate bifurcations, distinguishing successful reprogramming from dead-end trajectories (Fig. 3C). Notably, Ageas demonstrated superior predictive fidelity across all classification metrics (*Acc* = 0.736, *F*1_macro_ = 0.783, *AUROC* = 0.909), outperforming the second-best method, CoSpar (*Acc* = 0.691, *F*1_macro_ = 0.737, *AUROC* = 0.856) (Fig. 3D, Supplementary Fig. 3A, and Supplementary Table 2). These metrics underscore Ageas’ heightened sensitivity to fate-determinative chromatin features derived exclusively from cross-sectional data.

**Figure 3:**
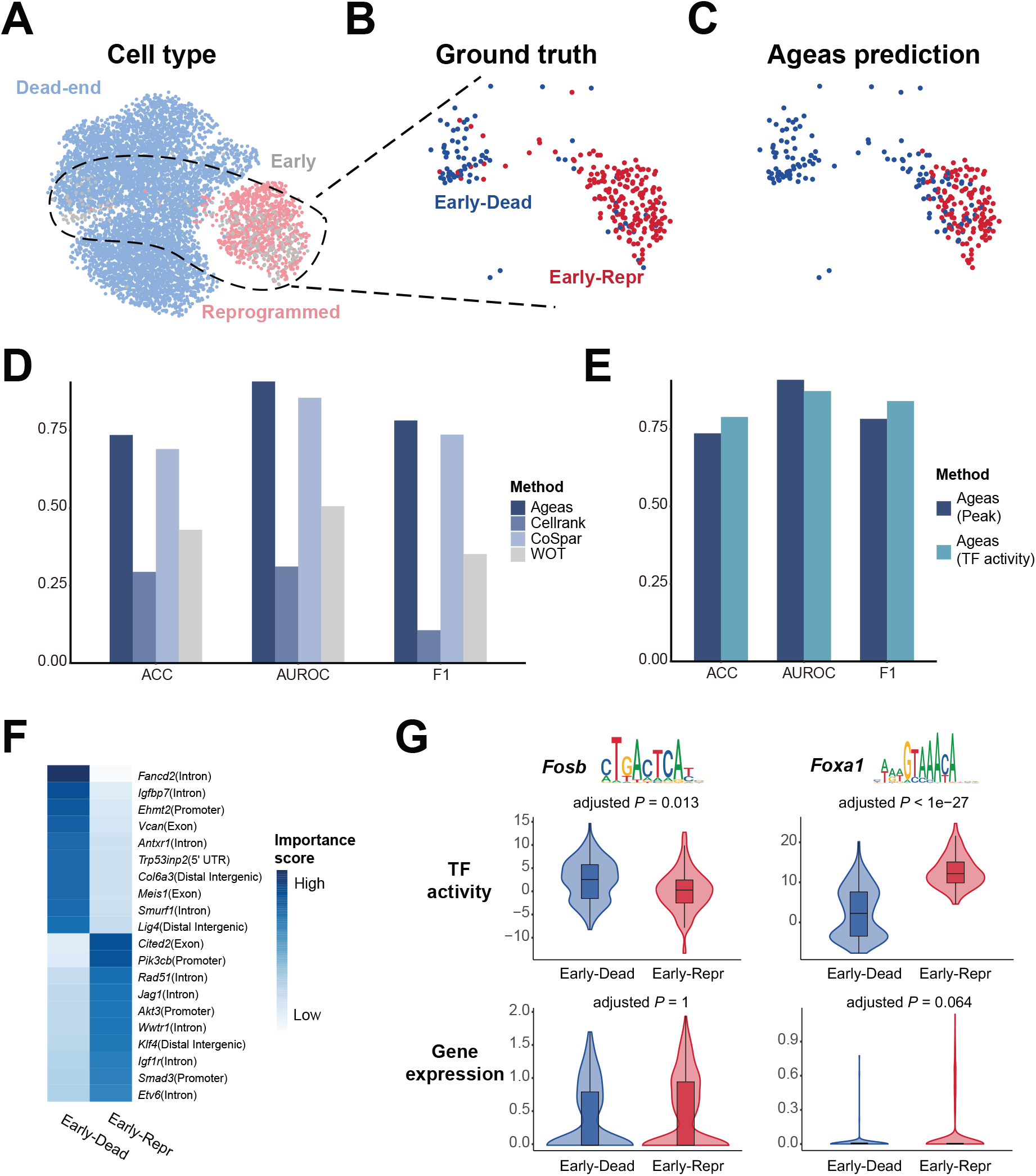
Ageas generalizes to epigenomic data to infer fate bias from chromatin accessibility. **A,** UMAP visualization of the CellTag-multi scATAC-seq reprogramming dataset, colored by cell type. **B,** Ground truth labels (*Early-Dead* vs. *Early-Reprogramming*) defined by lineage tracing. **C,** Ageas prediction of fate bias using only static chromatin accessibility data. **D,** Benchmarking of Ageas against CellRank, CoSpar, and WOT on scATAC-seq peak data, evaluated by accuracy, AUROC, and F1 score. **E,** Performance comparison of Ageas models trained on peak accessibility (Ageas-peak) versus ChromVAR-inferred transcription factor activity (Ageas-TF-activity). **F,** Heatmap showing the top fate-predictive chromatin peaks identified by the Ageas explanation module,annotated by their nearest genes and genomic region. **G,** Violin plots of inferred TF activity (*Fosb*, *Foxa1*) and gene expression in Early-Dead vs. Early-Repr cells, accompanied by their corresponding binding motif logos. Gene expression data are derived from the cell fates defined in Fig. 2C. Statistical significance was determined by the two-sided Wilcoxon rank-sum test with Bonferroni correction.

To systematically evaluate the influence of epigenomic feature representations on predictive fidelity, we benchmarked models trained on raw peak accessibility (Ageas-peak) against those utilizing ChromVAR-inferred transcription factor (TF) activities (Ageas-TF-activity). Ageas-peak achieved a higher AUROC (*AUROC*^peak^ = 0.909 vs *AUROC*^TF-activity^ = 0.872), whereas Ageas-TF-activity attained higher accuracy (*Acc*^peak^ = 0.736 vs *Acc*^TF-activity^ = 0.789) and 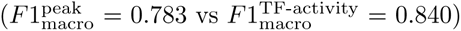 (Fig. 3E). We attribute this performance divergence to the intrinsic biological nature of the features: individual accessible loci capture early-stage regulatory priming, whereas aggregated TF activities reflect more stabilized, integrated transcriptional programs that consolidate during lineage commitment.

Leveraging the integrated explanation module, we interrogated the Ageas-peak model to identify specific chromatin loci driving fate predictions. Within the *Early-Dead* progenitor pool, predictive accessibility peaks were significantly enriched at loci associated with *Fancd2*, *Vcan*, and *Col6a3* (Fig. 3F). Those genes are linked to genome maintenance and fibroblast-associated extracellular matrix programs[50], constituting established molecular hallmarks of reprogramming resistance[51, 52]. Conversely, Ageas predicted that progenitors biased toward successful reprogramming based on the chromatin accessibility at specific regulatory elements, including those associated with *Jag1* and *Klf4*, both of which are established drivers of cell-state transitions[53, 54] (Fig. 3F). Furthermore, the most highly weighted fate-informative peaks mapped predominantly to intronic or distal enhancer elements rather than proximal promoters (Supplementary Fig. 4A). Consistent with previous studies, this indicates that enhancers are more closely associated with cell fate decisions than promoters[25, 55].

Subsequent interrogation of the Ageas-TF-activity model illuminated distinct regulatory networks segregating the two developmental trajectories. Crucially, numerous TFs exhibited profound divergence in inferred activity between *Early-Dead* and *Early-Reprogramming* states, despite lacking concomitant changes in their steady-state gene expression profiles. Specifically, *Early-Dead* cells were characterized by heightened activity of AP-1 complex member *Fosb*, and the homeobox factor *Hoxa9* (Fig. 3G, Supplementary Fig. 4B). These regulators are intimately linked to stress-responsive somatic maintenance[56] and the stabilization of refractory cellular identities[57]. In contrast, *Early-Reprogramming* cells demonstrated enriched activity of forkhead factors (*Foxa1* and *Foxd1*), consistent with chromatin priming and increased lineage plasticity during successful fate conversion[58] (Fig. 3G, Supplementary Fig. 4B). Collectively, these Ageas-driven multimodal analyses support the idea that epigenomic priming is associated with early fate divergence and can provide predictive information before overt transcriptional divergence.

### 2.5 Ageas reveals spatially coherent fate bias during axolotl spinal cord regeneration

To evaluate the efficacy of Ageas within a spatial transcriptomics framework, we analyzed a Stereo-seq dataset[59] profiling axolotl spinal cord regeneration at 15 days post-amputation, capturing an intermediate stage characterized by the resumption of active neurogenesis (Fig. 4A). Within this sample, two major regeneration lineage trajectories have been previously established: Lineage-1 which progresses from reactive ependymoglial cells (reaEGCs) through regenerative intermediate progenitor cells 1 (rIPC1) to yield immature motor neurons (IMN) and *nptx* -expressing excitatory neurons (nptxEX); and Lineage-2, which transits from reaEGCs through regenerative intermediate progenitor cells 2 (rIPC2) to specify dorsal progenitor excitatory neurons (dpEX) (Fig. 4A).

**Figure 4:**
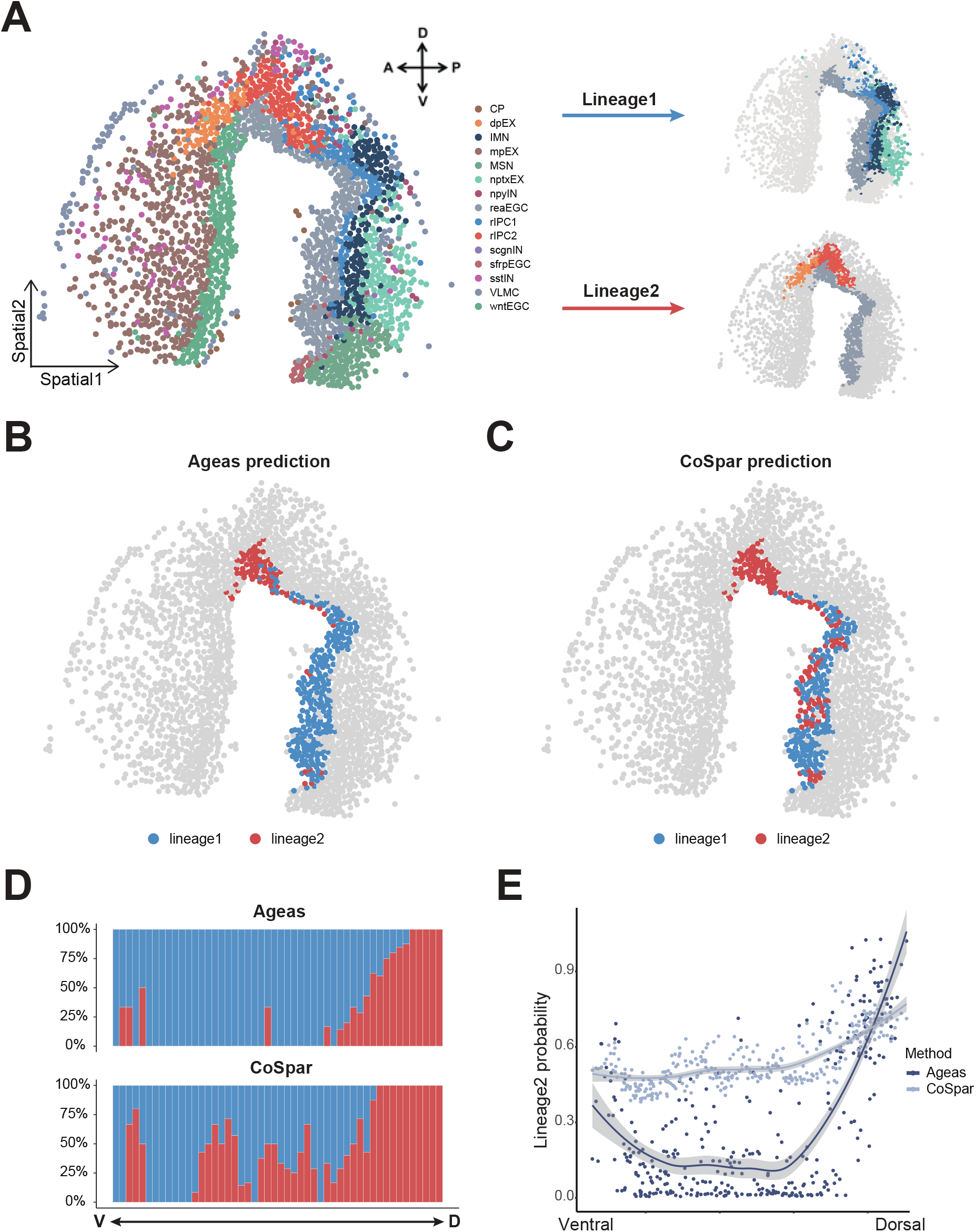
Ageas reveals spatially coherent fate bias during axolotl spinal cord regeneration. **A,** Spatial visualization of the regenerating axolotl spinal cord at 15 days post-amputation. Left: Cell type annotation. Right: Schematic of the two major regeneration lineages (Lineage-1 and Lineage-2) arising from reactive ependymoglial cells (reaEGCs). **B** Spatial visualization of predicted fate bias for reaEGCs toward Lineage-2 (red) or Lineage-1 (blue) using Ageas. **C** Spatial visualization of predicted fate bias using CoSpar. **D** Bar plots showing the predicted lineage composition of reaEGCs arranged along the V-D axis. **E** Scatter plot quantifying the predicted probability of Lineage-2 fate along the V-D axis. Curves represent the fitted trend for Ageas and CoSpar.

We tasked Ageas with resolving the bipartite fate bias of the reaEGC progenitor pool toward either Lineage-1 or Lineage-2 relying on static, time-agnostic transcriptomic profiles. To benchmark predictive performance against CoSpar, we designated reaEGCs as the progenitor population and the downstream regenerative states as terminal identities. Consequently, Ageas predicted a highly localized enrichment of Lineage-2-biased reaEGCs within the dorsal domain of the regenerating spinal cord (Fig. 4B,D). This predicted dorsal regionalization aligns well with previous reports which demonstrate that dorsal reaEGCs contribute to dpEX neurons during axolotl spinal cord regeneration[59]. In contrast, CoSpar generated spatially diffuse fate predictions that exhibited a less discernible dorsoventral (DV) organization (Fig. 4C,D). To rigorously quantify this spatial specificity, we mapped the Lineage-2 commitment probability along the DV axis. Ageas predictions gradually increased from low probabilities (0.0 ∼ 0.3) ventrally to high probabilities (*>* 0.8) dorsally, recapitulating the expected dorsal enrichment (Fig. 4E, Supplementary Fig. 5A). CoSpar predictions instead fluctuated around ∼ 0.5 across the axis, indicative of weak spatial resolution (Fig. 4E, Supplementary Fig. 5A).

Furthermore, we reasoned that progenitors sharing the same predicted fate should also exhibit spatial proximity, reflecting the localized microenvironmental coordination inherent to tissue regeneration[60–62]. To quantitatively assess this, we calculated the local same-type neighbor ratio, defined as the proportion of immediately adjacent cells sharing the same predicted lineage bias (Supplementary Fig. 5B). Ageas-predicted reaEGCs exhibited a significantly higher neighbor ratio than those predicted by CoSpar, confirming a pronounced local clustering of lineage-committed cells (Supplementary Fig. 5C).

Together, these findings demonstrate that Ageas can infer spatially coherent fate bias patterns from static spatial transcriptomic data, outperforming a representative time-aware method and revealing the intrinsic dorsal enrichment of neural regenerative potential in axolotl spinal cord repair.

### 2.6 Ageas infers spatially organized fate priming within the human epiblast

During human gastrulation, the epiblast serves as the pluripotent source for all three foundational germ layers: ectoderm forms directly from the epiblast, whereas mesoderm and endoderm emerge from epiblast cells that ingress through the primitive streak (PS)[63, 64]. In mouse, lineage tracing studies have revealed that the epiblasts exhibit lineage-specific biases, preferentially contributing to certain germ layers rather than uniformly giving rise to all lineages[65]. However, whether such fate bias exists in human epiblasts, and the mechanisms underlying this potential priming, remain largely unknown. To address this critical knowledge gap, we collected a 3D spatial transcriptomic atlas of a Carnegie stage 8 human embryo, generated by serial Stereo-seq sectioning[19]. This dataset reconstructs the entire embryo in three dimensions, comprising 62 consecutive slices aligned along the anterior-posterior (A-P) axis. Within this coordinate system, slices 1–2 define the anterior extreme, whereas slices 61–62 encompass the posterior pole (Fig. 5A,B, Supplementary Fig. 6). We applied Ageas to resolve the developmental trajectory of the epiblast toward primitive streak (PS) or ectoderm (Ecto) lineages, enabling us to investigate how spatial context and molecular programs contribute to early fate specification in human embryos.

**Figure 5:**
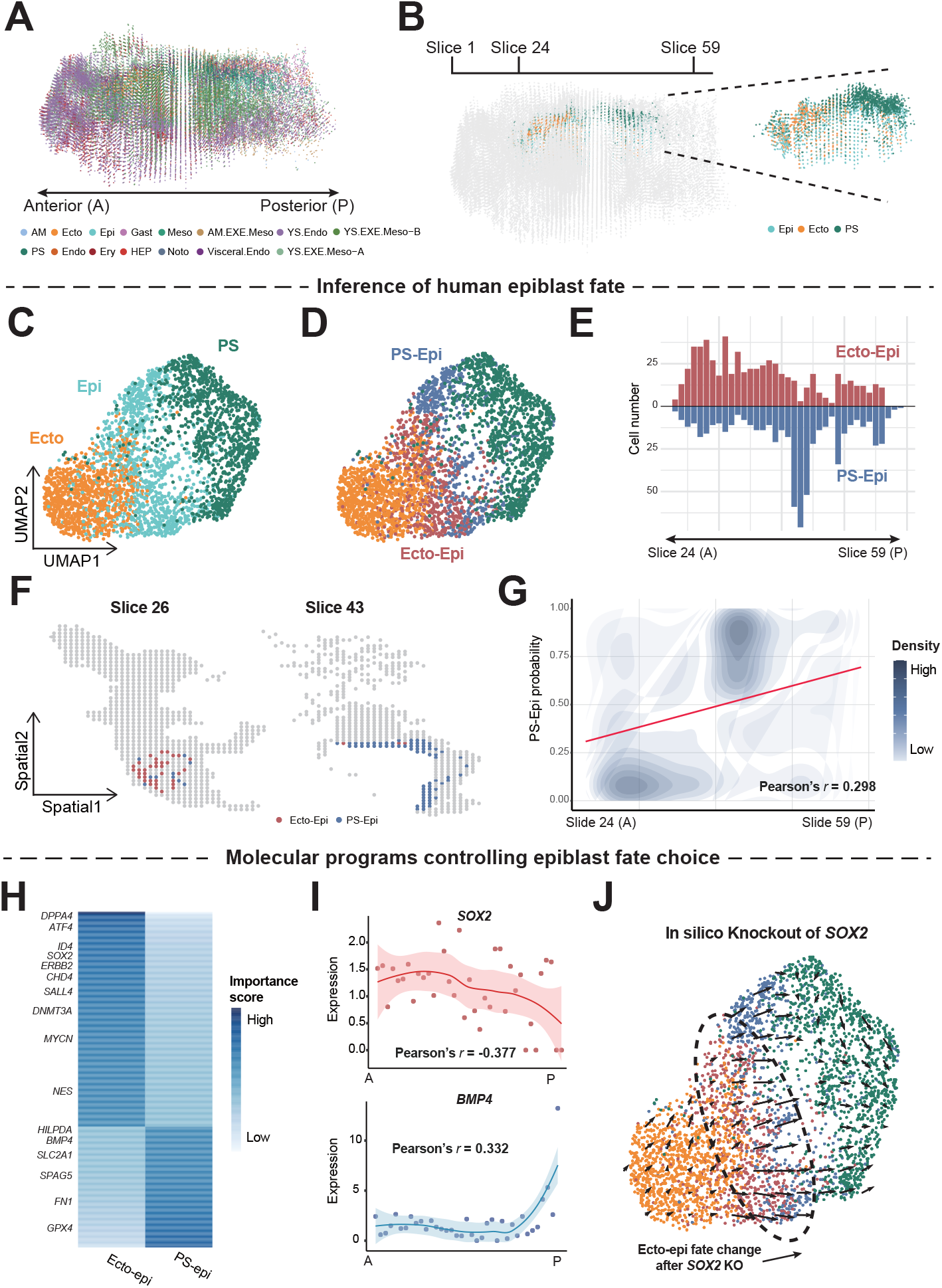
Spatially organized fate priming in the human epiblast. **A,** 3D reconstruction of a Carnegie Stage 8 human embryo profiled by Stereo-seq. Abbreviations: AM, amnion; Ecto, ectoderm; Epi, epiblast; Gast, gastrulating cells; Meso, mesoderm; AM.EXE.Meso, amniotic extra-embryonic mesoderm; YS.Endo, yolk sac endoderm; YS.EXE.Meso-A/B, yolk sac extra-embryonic mesoderm A/B; PS, primitive streak; Endo, endothelial; Ery, erythroid cells; HEP, hemogenic endothelial progenitors; Noto, notochord; Visceral.Endo, visceral endoderm. **B,** Representative spatial slices illustrating the selection of epiblast and downstream cells for fate prediction. Epiblast cells are present from Slice 24 to Slice 59 along the A-P axis. **C,** UMAPs of epiblast cells colored by celltype. **D,** UMAPs of epiblast cells colored by Ageas predicted fate. **E,** Ageas predicts epiblast fate specification (Ecto-Epi and PS-Epi) along the A-P axis. **F,** Spatial distribution of Ecto-Epi (blue) and PS-Epi (red) cells on representative slices. **G,** Correlation of predicted PS-Epi probabilities with the A-P spatial coordinate. Color represents the local density of epiblasts. For instance, high density in the upper-right region indicates that posterior epiblasts predominantly exhibit high primitive streak probability. **H,** Feature importance of top driver genes for Ecto-Epi and PS-Epi fates. **I,** Spatial expression trends of representative driver genes (*SOX2*, *BMP4*) along the A-P axis. Points represent mean expression of epiblast per slice; curves show the fitted trend with Pearson correlation coefficients. **J,** CellOracle in silico perturbation analysis simulating *SOX2* knockout. Vector field arrows indicate the predicted shift in cell state transitions, showing a loss of ectodermal priming trajectory.

Ageas was trained on transcriptomic profiles of the definitive terminal states (Ecto and PS) and then applied to predict each epiblast cell’s probability of adopting an ectodermal versus a PS-derived mesendodermal or endodermal fate. This analysis revealed two fate-biased epiblast populations, Ecto biased epiblast (Ecto-Epi) and PS biased epiblast (PS-Epi) (Fig. 5C,D). These findings are consistent with mouse lineage tracing studies[65], suggesting that spatially organized fate priming may be conserved between mouse and human epiblast development.

By quantifying the distribution of these fate-biased epiblast populations along the A-P axis, we identified a pronounced spatial pattern: Ecto-Epi were enriched on the anterior side, whereas PS-Epi were enriched on the posterior side (Fig. 5E,F). To quantify this predicted spatial gradient, we evaluated the correlation between the Ageas-predicted fate probability and the precise A-P coordinate. The probability of PS commitment increased progressively from anterior to posterior pole (Pearson’s *r* = 0.298*, P <* 2 × 10^−16^), concomitant with a symmetric decay in ectodermal bias (Pearson’s *r* = −0.298*, P <* 2 × 10^−16^) (Fig. 5G, Supplementary Fig. 7).

These findings indicate that human epiblasts exhibit regionally correlated fate potential, with anterior epiblasts preferentially committing toward ectodermal lineages and posterior epiblasts biased toward generating primitive streak-derived mesoderm and endoderm. Moreover, these results demonstrate that Ageas can reconstruct these spatial fate gradients from static transcriptomic data, offering a powerful framework for decoding human developmental fate decisions when direct experimental lineage tracing is practically prohibitive.

### 2.7 Ageas enables the virtual mechanistic interrogation of regional regulators driving epiblast fate

To explore the molecular programs regulating human epiblast fate decisions, we leveraged the integrated explainability module of Ageas to identify lineage driver genes defining the Ecto-Epi and PS-Epi populations. Based on the calculated feature importance scores, Ageas extracted 287 genes associated with Ecto-Epi bias and 161 genes associated with PS-Epi bias. The Ecto-Epi gene set was enriched for canonical regulators of pluripotency and neuroectodermal specification, including *SOX2* [66], *DPPA4* [67], and *ID4* [68], whereas the PS-Epi gene set featured well-known mesendodermal and epithelial-mesenchymal transition components, such as *BMP4* [69] and *FN1* [70] (Fig. 5H).

To examine whether these lineage drivers exhibit spatial patterning within the developing embryo, we evaluated the correlation between localized gene expression and anatomical positioning along the A-P axis. A total of 72 ectoderm-associated genes, including *SOX2*, *SALL4*, and *CEBPZ*, along with 24 primordial streak-associated genes, including *BMP4*, *CRABP2*, and *ID3*, demonstrated significant spatial correlation (|*r*| ≥ 0.2*, P <* 2 × 10^−16^) (Fig. 5I and Supplementary Fig. 8A). These results indicate that epiblast fate commitment is governed by regionally graded transcriptional programs, suggesting that early germ-layer patterning emerges from the spatial polarization of regulatory gene networks (GRNs) across the epiblast.

Given that *SOX2* and *CEBPZ* function as TFs, we hypothesized they play active, mechanistic roles in reinforcing this epiblast fate determination. To further explore the potential regulatory relevance, we deployed CellOracle[71], a GRN-based *in silico* perturbation framework, to simulate the targeted loss-of-function of these factors. The perturbation impaired the transition from epiblast to ectoderm predicted trajectories, suggesting that *SOX2* and *CEBPZ* actively maintain ectodermal fate priming (Fig. 5J and Supplementary Fig. 8B).

Collectively, these analyses demonstrate that Ageas-guided lineage driver discovery, coupled with *in silico* regulatory perturbation, enables virtual cellular interrogation of spatially organized fate priming in the human epiblast.

## 3 Discussion

Ageas introduces a classification-based strategy that learns latent “fate memory” features from single-cell or spatial data, eliminating any requirement for explicit time-course or pseudotime information. It captures the molecular memory that progenitor cells carry and uses this to predict their future fates. This approach contrasts with many trajectory methods that rely on temporal ordering, offering a more flexible way to infer cell fate purely from snapshot data[15, 16, 72, 73].

By employing an evaluation-and-selection-based ensemble architecture, Ageas achieves robust generalization across divergent data modalities. The framework systematically evaluates a diverse array of predictive algorithms, dynamically retaining only the highest-performing models for final integration. This adaptive design ensures the applicability of Ageas beyond standard single-cell transcriptomics, enabling high-fidelity predictions on other modalities such as chromatin accessibility. Ultimately, this modality-agnostic ensemble strategy future-proofs the analytical pipeline, ensuring its broad utility across rapidly evolving single-cell and spatial multi-omics technologies.

Ageas successfully tackles a challenging biological question of human epiblast fate priming that is difficult to interrogate using standard experimental lineage tracing, as such approaches rely on genetic manipulation that is not feasible in human embryos. While mouse lineage tracing has revealed the presence of fate bias within the epiblast, mechanistic interpretation is limited by the lag between labeling and sequencing: by the time clonal descendants are profiled, the original epiblast state and its spatial context are no longer directly observable[65]. Ageas circumvents this constraint by inferring fate bias directly from a static embryo snapshot and indicates that epiblast fate potential is spatially organized, linking lineage bias to anterior-posterior position. This observation is consistent with the established model and classical hypothesis wherein the precise spatial positioning of early embryonic cells exposes them to distinct signaling cues that drive specific lineage outcomes[19, 74–76]. The ability of Ageas to identify lineage-driving regulators makes it well suited for in silico perturbation studies. By prioritizing transcription factors and genes associated with fate decisions, Ageas can inform virtual cell frameworks that simulate cellular responses to genetic or chemical perturbations[77]. In this context, Ageas provides candidate regulators whose computational perturbation can be used to predict fate shifts, thereby facilitating hypothesis generation for experimental validation.

The architecture of Ageas is inherently flexible and can accommodate multiple data modalities for fate prediction. In particular, Ageas can be combined with multi-omics integration models that learn a shared latent representation across modalities, such as GLUE[78] or multiVI[79]. Applying Ageas to such integrated embeddings enables multimodal fate inference by leveraging complementary molecular information, thereby extending its applicability beyond single-modality analyses.

Ageas requires accurate annotation of terminal cell states for training and evaluation. If these endpoint labels are noisy or incorrect, the model’s learning and predictions will be compromised. In practice, this means Ageas is only as good as the reference fate definitions provided; assuming high-quality, confidently assigned terminal cell identities is essential for reliable results.

Taken together, Ageas provides a general, time-agnostic framework for cell fate inference that decouples fate prediction from temporal reconstruction, enabling robust cell fate analysis of single-cell and spatial multi-omics data.

## 4 Methods

### 4.1 The Ageas Framework

Ageas is a time-agnostic transfer-learning framework that infers progenitor fate bias by learning fate-discriminative molecular information from terminal cell states. Terminal cells with defined lineage identities are used as the reference population, whereas progenitor or intermediate cells constitute the query population whose future lineage outcomes are to be predicted. Both populations are represented in the same molecular feature space, allowing models trained on the terminal states to be directly applied to the query cells without requiring temporal ordering or trajectory reconstruction. To accommodate variation across datasets and molecular modalities, Ageas employs an AutoML-inspired strategy that evaluates candidate configurations spanning neural-network, linear, kernel-based, and tree-based classifiers. Models showing strong cross-validation performance on the terminal reference population are retained, retrained, and combined into a dataset-adaptive ensemble. The ensemble is subsequently applied to the progenitor population, and its averaged class scores are interpreted as relative fate biases toward the corresponding terminal outcomes. Ageas additionally integrates model-specific feature-attribution scores across the retained models to identify molecular features associated with each predicted fate.

#### 4.1.1 Candidate Neural-Network Models for Fate-Memory Learning

Ageas incorporates three neural-network families as candidate learners within its model-selection framework. These architectures provide complementary inductive biases for learning fate-discriminative patterns from terminal molecular profiles and are evaluated together with the classical machine-learning models described below. Importantly, the individual neural-network architectures are not assumed to be universally optimal; their inclusion in the final ensemble is determined by their validation performance on each dataset.

##### Neural Network Classification Pipeline

Let **x***_raw_* denote the initial input vectors. The data is first projected into an initial latent space via an embedding module:

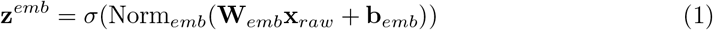

The resulting representation **z***^emb^* is sequentially processed through a series of cascaded structural stages, where each propagation through the *j*-th block of the *i*-th stage is formulated as:

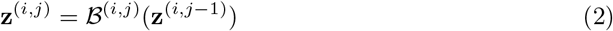

Here, *B*^(*i,j*)^ represents the specific building block employed, such as an MLP, ResNet Mixer, or RNN module, which are detailed in the subsequent sections.

Upon traversing all intermediate layers and aggregation steps, the final representation **z***_final_* is flattened into a single dense vector and subjected to a stochastic dropout regularization process. A terminal fully connected (*fc*) decision layer projects the regularized representation into the requisite logit space:

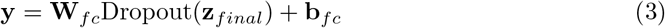

where **y** ∈ ℝ*^C^* represents the output logits over *C* target classes. The entire network framework is optimized end-to-end utilizing standard cross-entropy minimization across the training cohort.

##### Multilayer Perceptron (MLP) Architecture

We implemented a foundational feedforward module, comprising a linear transformation, normalization, and non-linear activation. Given an input tensor 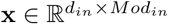, the forward propagation is formally defined as:

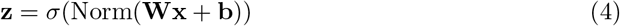

Here, 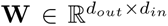 and 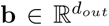 represent the learnable weight matrix and bias vector, respectively. The function Norm denotes the applied normalization strategy, such as layer normalization[80–82], while *σ*(*z*) = max(0*, z*) represents the element-wise Rectified Linear Unit (ReLU) activation[83]. The resulting tensor 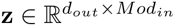 serves as the final representation for the module.

To facilitate robust feature extraction across varying dimensionalities while mitigating the vanishing gradient problem[84], we also constructed a residual-connected encoder module incorporating a parameter-efficient bottleneck architecture[84, 85]. This module introduces an intermediate latent dimensionality, 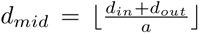, to gracefully map between the input dimension *d_in_* and the target output dimension *d_out_* by a factor of *a* ≥ 1 controlling the bottleneck compression ratio. The primary encoding pathway subjects the input 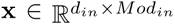 to a sequence of two cascaded linear transformations, projecting through an intermediate latent space before mapping to the target output dimension:

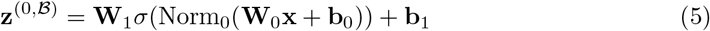

Here, 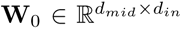 and 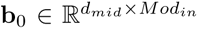 parameterize the initial projection into the bottleneck, while 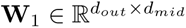 and 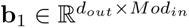 govern the subsequent expansion to the target dimensionality. In parallel, a residual connection *F_res_*(**x**) computes an identity mapping, or a linear projection shortcut if the input and output dimensionalities differ, ensuring dimensional compatibility for downstream aggregation:

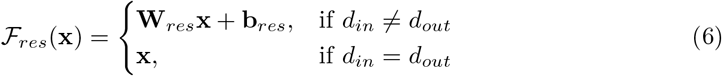

Finally, the outputs of the primary encoding and residual pathways are aggregated via element-wise addition, followed by terminal normalization and activation:

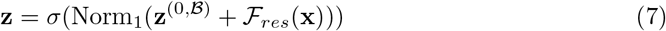

where 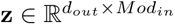 defines the final output tensor of the residual module.

##### Bidirectional Recurrent Neural Network (RNN) Architecture

To explicitly model complex, higher-order dependencies among genomic features, we extended our framework to incorporate bidirectional RNN modules[86, 87]. Although single-cell profiles lack an inherent temporal axis, processing the **z***^emb^* embedded latent features sequentially could allow the network to capture latent combinatorial gene interactions. To accommodate varying degrees of sequence complexity and memory requirements, the recurrent layer is dynamically configurable to utilize standard RNN[86, 87], LSTM[88], or GRU[89] transition functions implemented in PyTorch[90].

To capture hierarchical representations, each recurrent block *B* stacks *I* sequential layers. Let 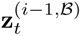 denote the input feature to the *i*-th internal layer at sequence index *t*, where the initial layer input 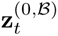 corresponds to the block input **x***_t_*. The bidirectional recurrent layer simultaneously processes the profile in both forward and reverse directions. At each sequence step, the chosen transition function *H* ∈ {RNN, LSTM, GRU} integrates the feature representation from the preceding layer with the recurrent memory from the adjacent sequence step:

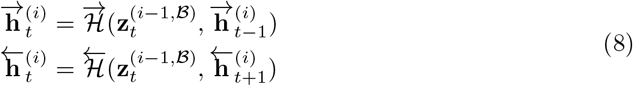

with initial hidden states 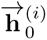 and 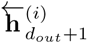 initialized to zero.

Notably, while standard RNN and GRU functions maintain only this hidden state 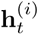, the LSTM configuration fundamentally relies on a dual-state representation, updating an internal memory cell state 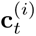 alongside the hidden state as an additional input and output of *H*, which is similarly initialized to zero at the sequence boundaries[88]. Regardless of the specific transition function type, the respective directional hidden representations are concatenated at each sequence index *t* to form the layer’s complete hidden state 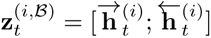 ensuring that the feature representation integrates global transcriptomic context. Thus, the final output of the *i*-th internal layer is:

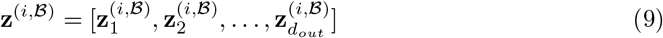

A key architectural novelty of Ageas is the adaptation of these temporal sequence processors for static omics data via a stabilized residual framework. To construct deeper architectures without suffering from degradation, we implemented a residual-connected block that wraps the entire multilayer recurrent transformation. Let **z**^(*I,B*)^ denote the complete output sequence generated by the terminal layer of the stack; the final normalized block output representation **z** aggregates this sequence with a parameterized shortcut connection:

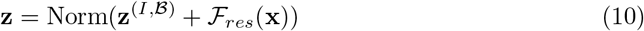

##### Factorized Global Spatial Mixing Architecture

Standard Convolutional Neural Networks (CNNs) are fundamentally designed to exploit local spatial invariances, which are typically absent in unstructured single-cell profiles like scRNA-seq gene expression matrices[91, 92]. To bridge the paradigm gap between structured convolutions and dense MLPs, we engineered a Mixer Classifier utilizing a ResNet[84]-based module configured explicitly for continuous global spatial mixing.

For an input tensor 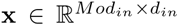, the network first projects the data into an expanded latent space via an initial global convolution without bias. The intermediate output is normalized, activated via a ReLU function (*σ*), and explicitly reshaped to an intermediate sequence length of 2*d_latent_* to restore spatial topology. A subsequent max-pooling operation, with a kernel size of 3, a stride of 2, and a padding of 1, halves this intermediate length, establishing the foundational feature sequence dimension *d_latent_* and channel dimension *Mod_latent_*:

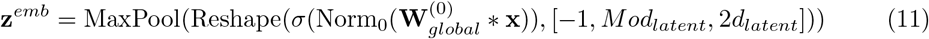

To explicitly model long-range, cross-gene dependencies without relying on a flattened, unstructured vector, our architecture employs a factorized residual bottleneck strategy.

Specifically, the mixer module isolates cross-channel interactions from spatial dependencies through a three-stage transformation sequence. First, an independent channel-mixing step is executed on the block input **x** using a 1 × 1 convolution without bias, projecting the representation into an intermediate bottleneck width *Mod_mid_*:

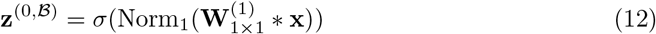

Second, spatial mixing is applied with the convolutional kernel size expandable to the current sequence length *d_in_*. This operation expands the features into a temporary composite dimension, acting analogously to the global receptive field of an MLP to synthesize systemic transcriptomic signatures:

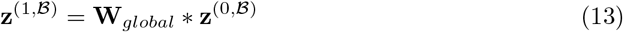

Crucially, to preserve the dimensional topology required for deep sequential processing, the output **z**^(1*,B*)^ is dynamically reshaped to reinstate the sequence length *d_out_*determined by the target spatial stride, before applying normalization and activation:

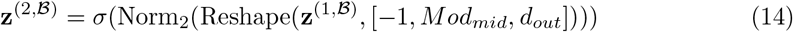

Third, a final 1 × 1 convolutional channel-refinement step expands the bottleneck to the target output dimensionality *Mod_out_*:

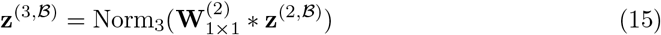

The module concludes by integrating these factorized transformations with the original identity mapping **x** via a skip connection, incorporating a convolutional downsampling operation *F_downsample_*if the spatial or channel dimensionalities differ:

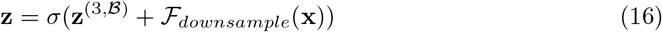

Finally, unlike standard MLPs that directly flatten intermediate representations, our sequence-aware Mixer Classifier aggregates the globally mixed feature maps using 1D Adaptive Average Pooling. This operation distills the comprehensive global sequence context into a fixed-length spatial embedding prior to terminal classification, ensuring robust and translation-invariant fate inference.

##### Gradient-based Interpretability Framework

To extract key regulatory features driving fate decisions, Ageas employs Integrated Gradients[93, 94] (IG) to compute feature attributions by integrating model gradients from a defined baseline. We initialize this baseline as a zero-tensor, representing a theoretically “silent” cellular state. For an input profile *x* assigned to class *c*, the vector **IG**(*x, c*) quantifies each feature’s local predictive contribution. To derive global regulatory signatures, we average these local attributions across all *N* cells predicted as class *c*, yielding a mean attribution vector ***µ****_c_*:

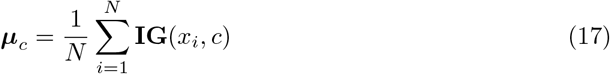

Applying *L*1 normalization standardizes these values across diverse modalities, producing a relative importance score **s***_c_*that represents proportional feature contributions:

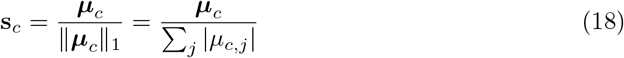

While **s***_c_* highlights salient features, it may conflate unique lineage drivers with general regulators active across multiple lineages. To isolate class-specific regulators, we calculate a discriminative specificity score **d***_c_*. This metric penalizes broad signals by subtracting the maximum importance observed in any background class k ≠ c from its target class importance:

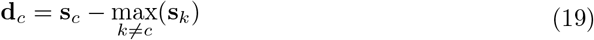

Analogous to an *in silico* differential analysis, this formulation amplifies regulatory elements uniquely predictive of a specific lineage commitment while explicitly suppressing generalized background noise.

#### 4.1.2 Classical Machine Learning and Tree-Based Architectures

Besides the deep learning architectures described in Section 4.1.1, we additionally integrated a robust suite of classical and tree-based algorithms. These established paradigms are encapsulated within the same centralized orchestration module, ensuring a mathematically rigorous and unified evaluation against their deep learning counterparts.

##### Logistic Regression

To establish a robust linear baseline and evaluate the linear separability of the cellular samples, we implemented a multinomial logistic regression model with scikit-learn[95]. This architecture optimizes a multi-class softmax objective, projecting the input features into a probabilistic class space via a linear transformation. The model is optimized using standard quasi-Newton solvers with configurable *ℓ*_1_ or *ℓ*_2_ regularization to induce feature sparsity and prevent overfitting. Crucially, this architecture is inherently interpretable, as feature importance can be directly inferred from the magnitude of the learned weight coefficients. To integrate these feature importance scores with those derived from deep learning models, we apply *L*1 normalization to the target class coefficient vector **w***_c_*:

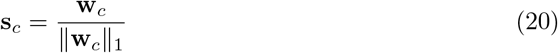

The normalized importance score **s***_c_* is then subjected to the same discriminative specificity calculation (**d***_c_*) defined in Equation 19.

##### Support Vector Machine

To capture distinct decision boundaries, we incorporated Support Vector Machine (SVM) classifiers implemented via the scikit-learn framework[95]. While logistic regression optimizes for probabilistic class assignment, SVM fundamentally differs by seeking to maximize the geometric margin between divergent cellular trajectories. Although the Ageas framework supports multiple kernel projections, we predominantly utilize the linear kernel when model explainability is required. Restricting the SVM to a linear kernel ensures that, similar to the logistic regression approach, the relative importance of specific molecular features can be directly extracted from the orthogonal vector defining the maximum-margin hyperplane. These extracted coefficients are subsequently normalized via Equation 20 to allow for direct comparison within the unified Ageas interpretability framework.

##### Tree-Based Ensembles

To resolve highly non-linear fate decisions and complex gene-gene interactions without requiring explicit kernel mappings, we integrated gradient-boosted decision trees using the XGBoost framework[96]. Unlike linear baselines, tree-based models dynamically partition the feature space, making them highly adept at handling heterogeneous single-cell modalities. To maintain the biological interpretability required by Ageas, the XGBoost module is mathematically coupled with SHapley Additive exPlanations[97] (SHAP). Specifically, we utilize the TreeExplainer module to process the entire subset of cells assigned to target class *c* simultaneously. This operation directly yields a unified, class-specific measure of feature attribution ***µ****_c_* without requiring single-cell level averaging:

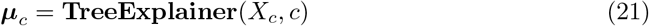

Here, *X_c_*denotes the matrix of all cellular profiles predicted as class *c*. Following this extraction, the feature importance scores are processed identically to the Integrated Gradients output from the deep learning models. The attribution vector ***µ****_c_* is standardized via *L*1 normalization to compute **s***_c_* (Equation 18), and subsequently refined via the discriminative specificity function **d***_c_* (Equation 19) to isolate unique lineage drivers.

#### 4.1.3 Data-Adaptive Model Selection and Ensemble Construction

To transition from isolated model training to an orchestrated, robust ensemble, we engineered an automated model selection pipeline that dynamically evaluates a diverse algorithmic spectrum. This module attempts to minimize manual hyperparameter selection biases, systematically curating a robust computational cohort for the final biological consensus.

The selection process relies on a rigorous multi-iteration *K*-fold cross-validation scheme to evaluate model configurations before instantiating the final ensemble. Initially, the centralized repository provides a vast array of candidate model configurations, denoted as the set *M*, encompassing varied architectures, hyperparameters, and optimization schedules. During each evaluation deployment, the single-cell profiles are stratified into training, validation, and test cohorts, preserving the inherent class distribution of rare cellular subpopulations. For each candidate configuration *m* ∈ *M*, the model *m_k_* is instantiated and trained on the training cohort of the *k*-th validation fold. Correspondingly, the performance of each trained model instance is evaluated on the held-out validation fold with a chosen evaluation metric, for example of *Acc* or *F*1_macro_, yielding a performance score *S_m__k_* for each fold *k*. The overall viability of the model configuration is then calculated as the mean score across all *K* trained instances:

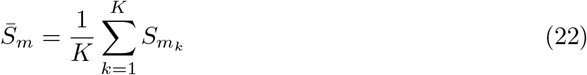

To prune underperforming blueprints while retaining a robust ensemble, model configurations are ranked based on their aggregated performance *S̅_m_* such that the highest-scoring configuration receives a rank *r*(*m*) = 1. We define an expected survival count *E* based on a predefined selection ratio *ρ*:

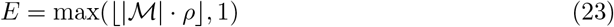

Besides this relative ranking condition, Ageas also enforces an absolute performance threshold to isolate potentially high-performing model configurations. Let *τ*_cut_ represent the minimal acceptable cutoff threshold, and *τ*_ret_ denote the high-performance retention threshold. A model configuration *m* is retained in the updated survivor cohort M′ if and only if it satisfies the following logical condition:

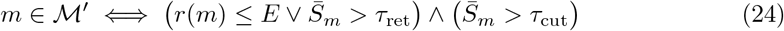

This dual-gate mechanism aggressively prunes the field of configurations while guaranteeing that highly specific, top-performing blueprints survive the selection process regardless of the stringent pruning ratio.

Following the iterative pruning, the surviving model configurations in *M*′ are utilized for a final training. For each surviving configuration, a new model instance is initialized and fully trained on the primary dataset. Trained models that exceed the rigorous retention threshold *τ*_ret_ during this final evaluation phase are directly aggregated into the terminal predictive ensemble. To generate robust fate predictions, this final ensemble mathematically averages the diverse probabilistic outputs from its constituent trained models.

#### 4.1.4 Ensemble-Based Identification of Fate-Associated Features

Beyond predicting progenitor fate bias, Ageas aggregates feature-attribution scores across the retained models to identify molecular features consistently associated with each lineage outcome. Because different model families capture complementary linear and nonlinear predictive patterns, their integrated attributions can provide a broader representation of fate-associated molecular signals than any single model. Ageas further implements an iterative feature-masking procedure to reduce the dominance of highly predictive markers and nominate additional, weaker signals associated with fate discrimination.

##### Performance-Weighted Ensemble Aggregation

During the model explaining phase, the framework harmonizes the specificity scores **d***_m,c_* derived from each surviving model *m* for a target class *c*. To ensure that highly accurate models contribute more heavily to the biological consensus than borderline models, we scale each model’s attribution vector by its cross-validated performance metric *S̅_m_*. The integrated ensemble attribution score **A***_c_* is computed as the sum of these performance-weighted vectors:

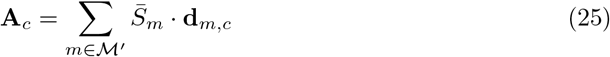

##### Iterative Upper-Outlier Masking

To uncover subtle yet important regulatory drivers that might be eclipsed by dominant markers, the framework employs an *N* -iteration sequential extraction algorithm. In each iteration *i*, the ensemble computes the integrated attribution scores 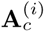. Ageas then identifies hyper-dominant features by calculating the Interquartile Range (IQR) of the importance distribution. Features whose scores exceed the upper outlier threshold, defined as *τ*_outlier_ = *Q*_3_ + *α* · IQR, where *α* is a configurable distance multiplier, are recorded as primary drivers and subsequently explicitly masked from the input dataset. The entire configuration selection and model training pipeline is then iteratively executed on this restricted feature space. By iteratively stripping away the most obvious signals, the models are forced to identify subtle regulatory modules governing the lineage bifurcation, ultimately approximating a comprehensive, multi-tiered hierarchy of cell fate drivers.

### 4.2 Datasets used in this study

To evaluate the Ageas framework, we employed a diverse collection of datasets, including scRNA-seq, scATAC-seq, and spatial transcriptomics.

#### 4.2.1 Classification benchmark datasets for fate memory learning

**Zheng68k**[30] is a classic peripheral blood mononuclear cell (PBMC) dataset generated via 10X Genomics, widely utilized for benchmarking cell type annotation performance. It contains approximately 68, 450 cells encompassing 11 distinct subtypes.

**Fu**[31] contains approximately 9, 000 human PBMCs sequenced using 10X Genomics technology. It provides a diverse and heterogeneous profile of immune cell states from clinical samples.

**MacParland**[32] consists of approximately 8, 444 cells from human liver tissue sequenced using the 10X Genomics Chromium platform. It encompasses 20 distinct cell populations, including hepatocytes, hepatic stellate cells, and a diverse landscape of liver-resident immune cells such as Macrophages, NK cells, and T cells.

**Zhang**[33] comprises approximately 12, 339 cells from human lung cancer and adjacent normal tissues sequenced via 10X Genomics. It covers various epithelial and immune cell types, reflecting the transcriptomic landscape of the tumor microenvironment.

#### 4.2.2 Benchmark datasets with lineage-tracing ground truth

**Mouse Hematopoiesis Dataset**[6] profiles the differentiation of hematopoietic stem and progenitor cells (HSPCs) into downstream lineages. From the original data, we selected a subset of 49, 116 cells possessing valid clonal barcodes for further analysis. This dataset establishes ground truth fate bias using LARRY (Lineage and RNA recovery), a lentiviral barcoding library. By labeling early progenitors, this method enables the retrospective identification of cell fate based on the lineage outcomes of their clonal siblings found in differentiated states.

**IEP Reprogramming Dataset**[25] captures the reprogramming of mouse fibroblasts into successfully reprogrammed iEPs or dead-end states. From the original data, we selected a subset of 8, 137 cells possessing valid clonal barcodes. This dataset utilizes CellTag-multi, a combinatorial lentiviral barcoding strategy, to establish ground truth by tracking clonal families from Day 3 progenitors to their terminal Day 28 fate outcomes.

#### 4.2.3 Epigenomic benchmark datasets

The iEP reprogramming scATAC-seq dataset profiles chromatin accessibility during the fibroblast reprogramming process. From the original experimental data, we selected a subset of 6, 799 cells possessing valid clonal barcodes. This dataset leverages the same CellTag-multi barcoding system[25] to provide ground truth labels, enabling the prediction of early fate bias directly from epigenomic snapshots. To evaluate the impact of different epigenomic feature representations on fate prediction, we utilized chromVAR[98] to calculate single-cell TF activity based on the cisBP motif database[99], facilitating a comparative analysis between integrated TF activity and raw peak accessibility profiles.

#### 4.2.4 Axolotl spinal cord regeneration spatial transcriptome

The axolotl spinal cord regeneration dataset profiles the transcriptomic landscape at Day 15 post-injury via Stereo-seq, capturing an intermediate stage of active neurogenesis[59]. Comprising 2, 396 spots across 15 cell types, this dataset resolves the spatial differentiation trajectories of reaEGCs, serving as a basis for inferring progenitor fate bias within the regenerative microenvironment.

#### 4.2.5 Human embryo 3D spatial transcriptomic atlas

The human embryo 3D spatial transcriptomic atlas comprises a CS8 embryo reconstructed from 62 serial Stereo-seq sections aligned along the anterior-posterior axis[19]. This dataset encompasses 38, 562 spatially resolved spots annotated across 15 distinct cell types. We leveraged this atlas to resolve the early fate priming of epiblast during gastrulation, enabling the characterization of regionally graded fate bias toward ectodermal and primitive streak-derived lineages.

#### 4.2.6 Data processing and feature selection

Single-cell and spatial transcriptomic data were processed using Scanpy[100]. Gene expression matrices were normalized to a library size of 10, 000 and log-transformed. We selected the top 2, 000 highly variable genes for downstream classification and fate inference analyses. For scATAC-seq data, we utilized Signac[101] for preprocessing. The peak-by-cell matrix underwent TF-IDF normalization, and the top 50, 000 accessible peaks were selected based on total counts for downstream analysis.

### 4.3 Methods included in benchmarking

#### 4.3.1 Cell classification methods

To assess the foundational ability of Ageas to learn molecular features from labeled data, we compared its performance against four representative cell type annotation tools.

**Seurat**[34] utilizes an anchor-based integration strategy to map query cells onto a reference manifold by identifying mutual nearest neighbors that represent shared biological states. To transfer cell type labels, we projected the query data onto the reference PCA structure and classified cells using a weighted vote based on anchors identified within the first 30 principal components.

**SingleR**[35] performs reference-based annotation by computing the Spearman rank correlation between query single-cell expression profiles and reference bulk or single-cell transcriptomes. We assigned labels based on the 80th percentile of correlation values across reference samples and applied iterative fine-tuning to resolve ambiguities among closely related cell subtypes using discriminative variable genes.

**scANVI**[36] is a deep generative model that extends variational autoencoders to perform semi-supervised learning, mapping both labeled and unlabeled cells into a shared latent embedding corrected for batch effects. We trained the model using a 10-dimensional latent space and two hidden layers with 128 units each, utilizing a zero-inflated negative binomial likelihood to model the count data distribution.

**TOSICA**[37] employs a Transformer-based architecture with self-attention mechanisms to capture global gene-gene interactions and project cells into an interpretable space without prior dimensionality reduction. We configured the model with a standard multi-head attention structure (8 heads) and a depth of 2 layers to extract context-aware molecular features and map query cells to reference cell types.

#### 4.3.2 Fate inference methods

To evaluate the accuracy of progenitor fate prediction, we benchmarked Ageas against three fate inference frameworks that rely on different computational paradigms. To ensure a fair comparison, we withheld real-time information from all methods. Instead, we uniformly assigned an “early” time label to progenitor cells and a “late” label to progeny cells.

**WOT**[13] uses optimal transport to infer cell state transitions across time points while accounting for cell proliferation and death. We reconstructed temporal couplings using entropic regularization (epsilon = 0.05) to account for stochasticity and applied asymmetric marginal constraints (lambda1 = 1, lambda2 = 50) to accommodate variations in cellular proliferation and growth rates.

**CoSpar**[14] optimizes the inferred transition matrix based on coherence and sparsity principles to robustly predict cell fate. We utilized the infer_Tmap_from_state_info_alone mode to calculate the transition matrix solely from transcriptomic profiles, applying multiscale smoothing (smooth_array = [20, 15, 10]) to enforce local coherence and a sparsity threshold (sparsity_threshold = 0.2) to prune low-probability transitions.

**CellRank**[16] models cellular dynamics as a Markov chain by coupling cell states across experimental time points using the RealTimeKernel. We inferred temporal couplings by solving a TemporalProblem via moscot to combine intra- and inter-time point transitions, enabling the computation of absorption probabilities toward terminal macrostates.

#### 4.3.3 Evaluation Metrics

To assess the robustness of cell type classification, we implemented a five-fold cross-validation strategy, iteratively training on 80% of the data and validating on the remaining 20%. For cell fate inference tasks, performance was quantified using Accuracy, F1 score, and AUROC. To ensure consistent AUROC calculation across different frameworks, predicted probabilities from all methods were scaled to the [0, 1] range, ensuring that the probabilities for binary fate outcomes summed to unity. The local same-type neighbor ratio was calculated to quantify the spatial coherence of fate predictions. For each spot, we identified the k=8 nearest spatial neighbors based on Euclidean distance. We then computed the proportion of these neighbors that shared the identical predicted fate label. Finally, a global coherence score was derived by averaging these local ratios across all spots.

#### 4.3.4 Peak annotation

We used the R package ChIPseeker[102] to annotate the scATAC-seq peaks from the CellTag-multi dataset. Peaks were annotated relative to the reference genome (mm10) to identify genomic locations, categorizing them as promoter, intron, exon, or distal intergenic regions.

#### 4.3.5 In silico perturbation analysis

We performed GRN-based in silico perturbation using CellOracle[71] (v0.10.0). A baseGRN was constructed using public scATAC-seq data[103] by identifying transcription factor (TF) binding motifs within accessible chromatin regions. This prior network (baseGRN) was integrated with our scRNA-seq data, and a Ridge regression model was fitted to estimate regulatory probabilities between TFs and targets. To ensure robustness, the inferred GRN was pruned to retain only statistically significant connections (*P <* 0.001) and the top 2,000 edges based on regulatory weight. Finally, we simulated TF knockouts by setting their expression to zero and using CellOracle’s signal propagation model to predict resulting shifts in cell fate trajectories.

## Supporting information

Supplementary Table 1. Five-fold cross-validation performance of Ageas and benchmark cell-type annotation methods across four scRNA-seq datasets.

Supplementary Table 2. Performance of Ageas and benchmark fate-inference methods on transcriptomic and chromatin-accessibility lineage-tracing dataset

Supplementary Table 3. Fate-bias scores and spatial correlations for candidate regulatory genes associated with ectodermal and primitive streak primin

## 5 Acknowledgements

We would like to express our sincere gratitude to Lei Hu and Ruodi Yang from Westlake University for their insightful discussions and valuable feedback on this work. We are also grateful to Jie Wang at the Guangzhou Institutes of Biomedicine and Health (GIBH), Chinese Academy of Sciences, for the inspiring discussions, and to Masayoshi Nakamoto for generous technical support throughout the project. We further thank Dr. Timothy Ramnarine from the Gene Center and Department of Biochemistry, Ludwig-Maximilians-Universität München, for his careful proofreading of the manuscript.

## Declarations

### Funding

This work was carried out without external funding support. All datasets used in this study are publicly available, and all analyses were performed using personal computational resources of the authors.

### Competing interests

All authors declare no competing interests.

### Data availability

The datasets utilized in this study are available at the following repositories: the Zheng68k PBMC dataset is available at SRA under accession SRP073767; the Fu PBMC dataset is available at Zenodo (https://zenodo.org/records/10947879); the MacParland human liver dataset is available at GEO under accession GSE115469; the Zhang lung cancer dataset is available at GEO under accession GSE146771; the mouse hematopoiesis (LARRY) lineage-tracing dataset is available at GEO under accession GSE140802; the iEP reprogramming (CellTag-multi) scRNA-seq and scATAC-seq datasets are available at GEO under accession GSE216521; the axolotl spinal cord regeneration Stereo-seq dataset is available at CNGBdb under accession CNP0002068; and the human embryo 3D spatial transcriptomic atlas is available at GSA under accession HRA005567.

### Code availability

The source of Ageas is available on Github (github.com/MaftyLab/Ageas), and the documentation on GitHub Pages including tutorial notebooks.

### Author Contributions

J.J. and G.Y. conceived the project, developed the Ageas framework, conducted all data analyses, and wrote the manuscript. A.K. provided advice on the data analysis and critically reviewed the manuscript. All authors read and approved the final manuscript.

## Supplementary Figures

**Supplementary Fig. 1:**
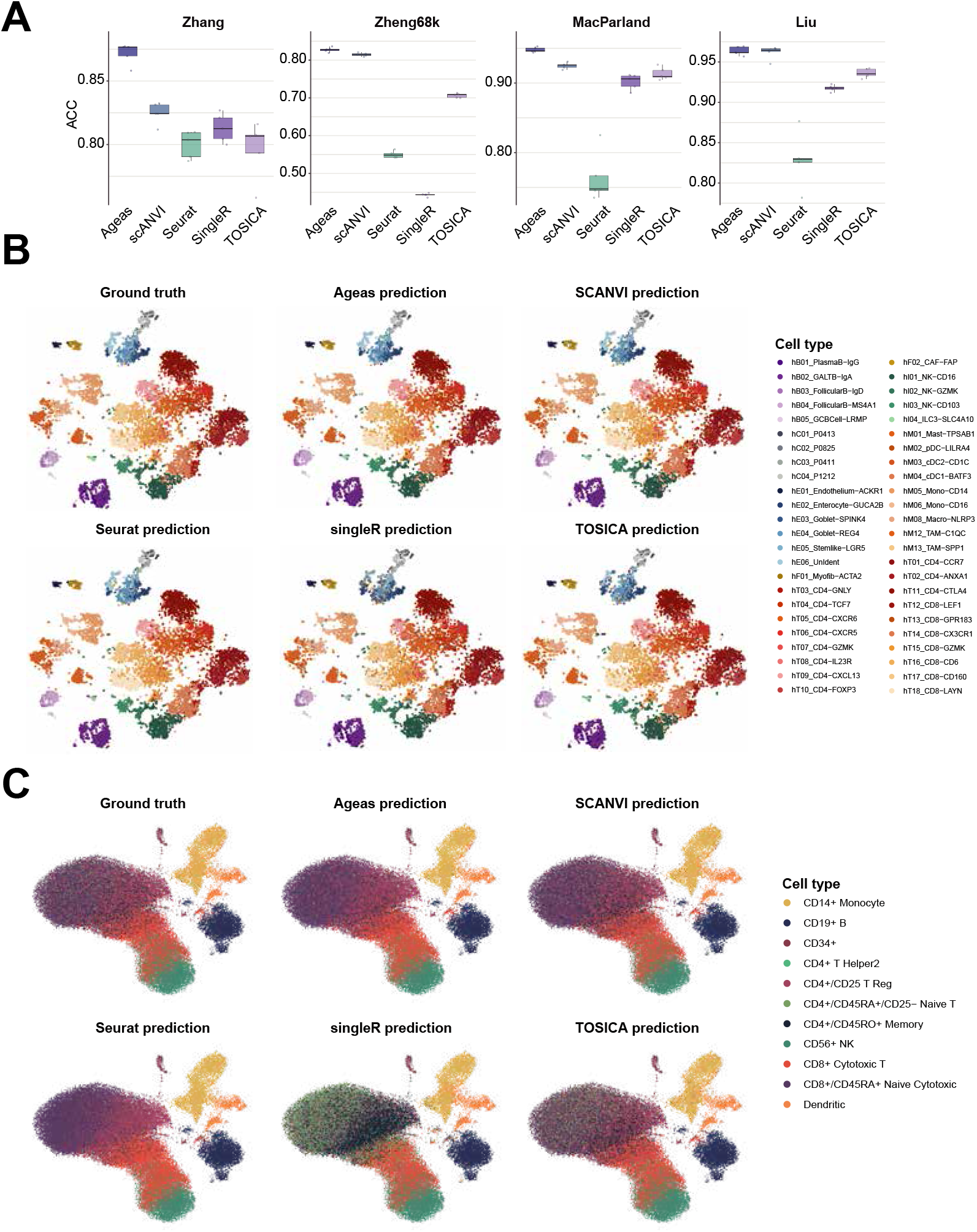
Classification performance benchmark of Ageas. **A,** Five-fold cross-validation accuracy of Ageas compared to four existing methods (scANVI, Seurat, SingleR, TOSICA) across four scRNA-seq datasets. Data are presented as mean ± s.d. **B-C,** UMAP visualizations comparing ground truth cell type labels with predictions generated by Ageas and benchmark methods for the Liu colon (B) and Zheng68k (C) datasets.

**Supplementary Fig. 2:**
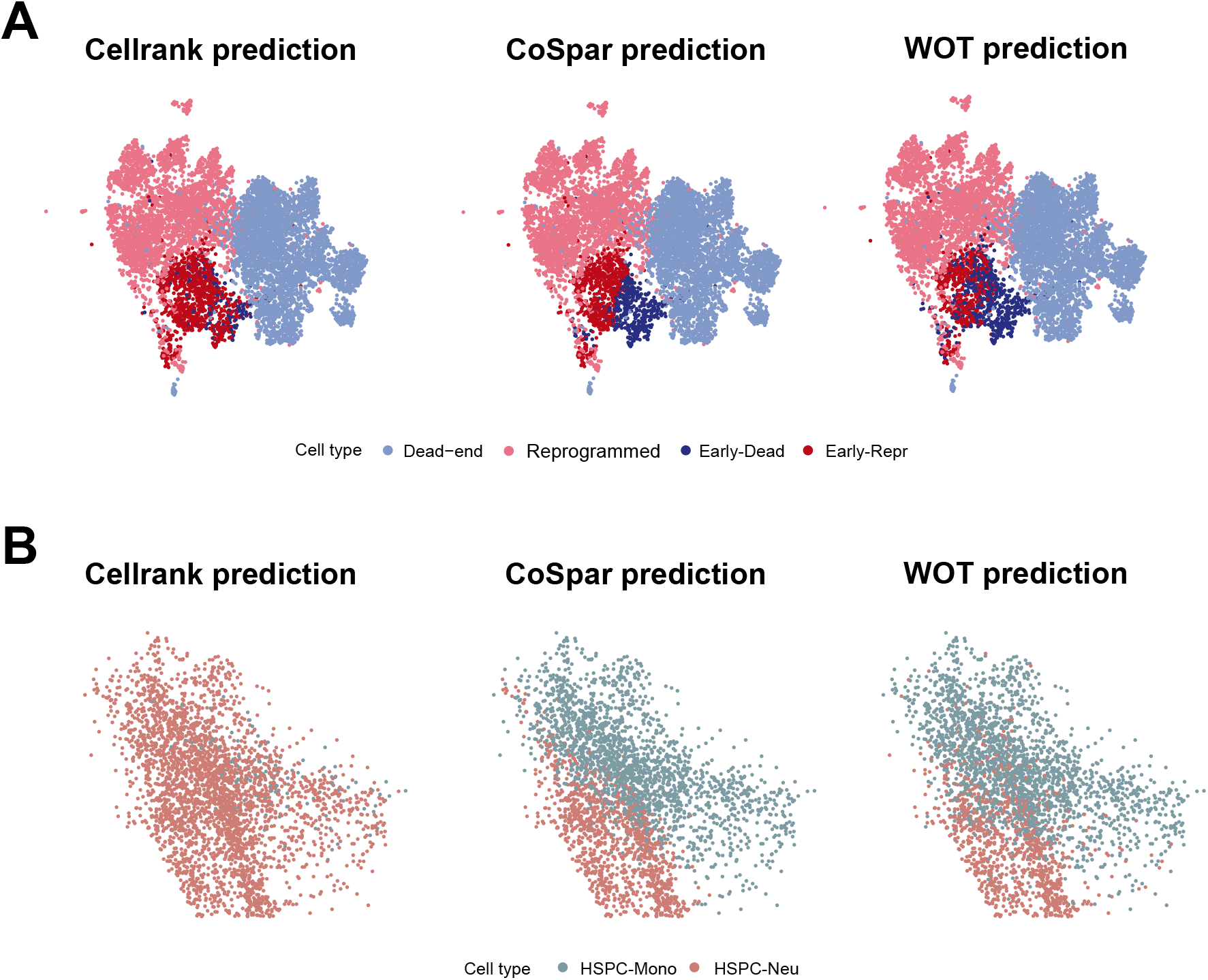
Fate prediction results for lineage-traced scRNA-seq datasets. **A-B,** Predicted cell fate labels of CellRank, CoSpar, and WOT for the iEP reprogramming (A) and mouse hematopoiesis (B) datasets.

**Supplementary Fig. 3:**
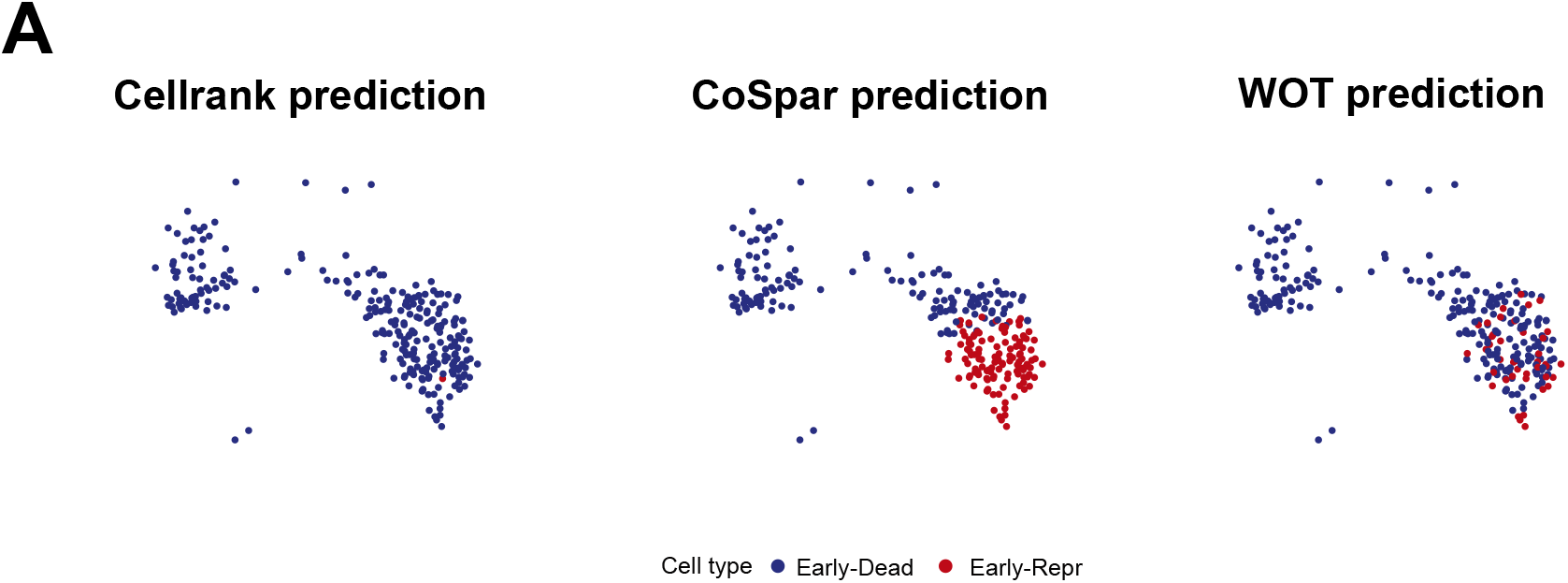
Fate prediction results for lineage-traced scATAC-seq dataset. **A,** Predicted cell fate labels of CellRank, CoSpar, and WOT for the scATAC-seq dataset.

**Supplementary Fig. 4:**
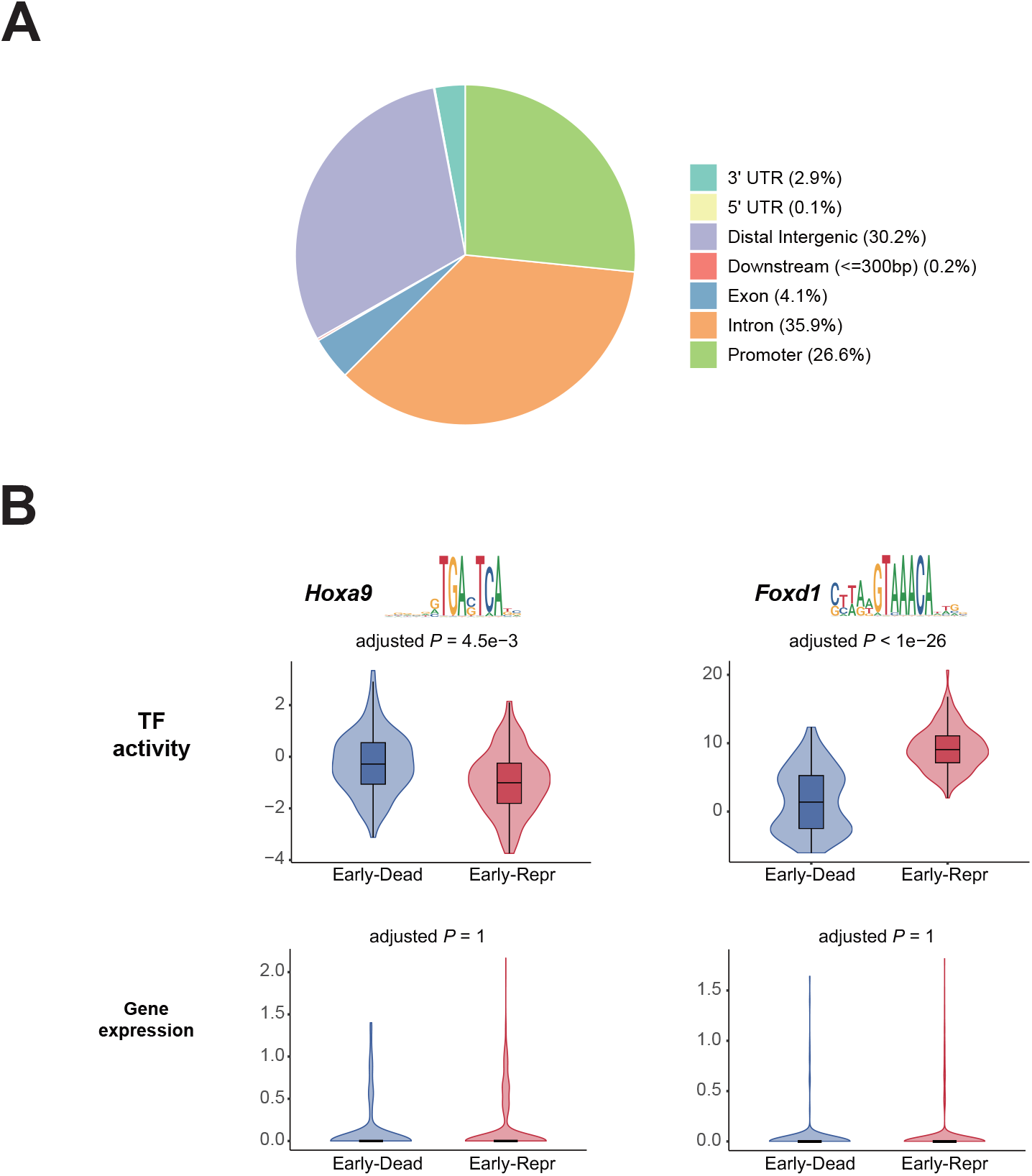
Genomic features of epigenomic fate drivers. **A,** Genomic annotation of top fate-informative peaks identified by Ageas explanation module. **B,** Violin plots of inferred TF activity (*Hoxa9*, *Foxd1*) and gene expression in Early-Dead vs. Early-Repr cells, accompanied by their corresponding binding motif logos. Statistical significance was determined by the two-sided Wilcoxon rank-sum test with Bonferroni correction.

**Supplementary Fig. 5:**
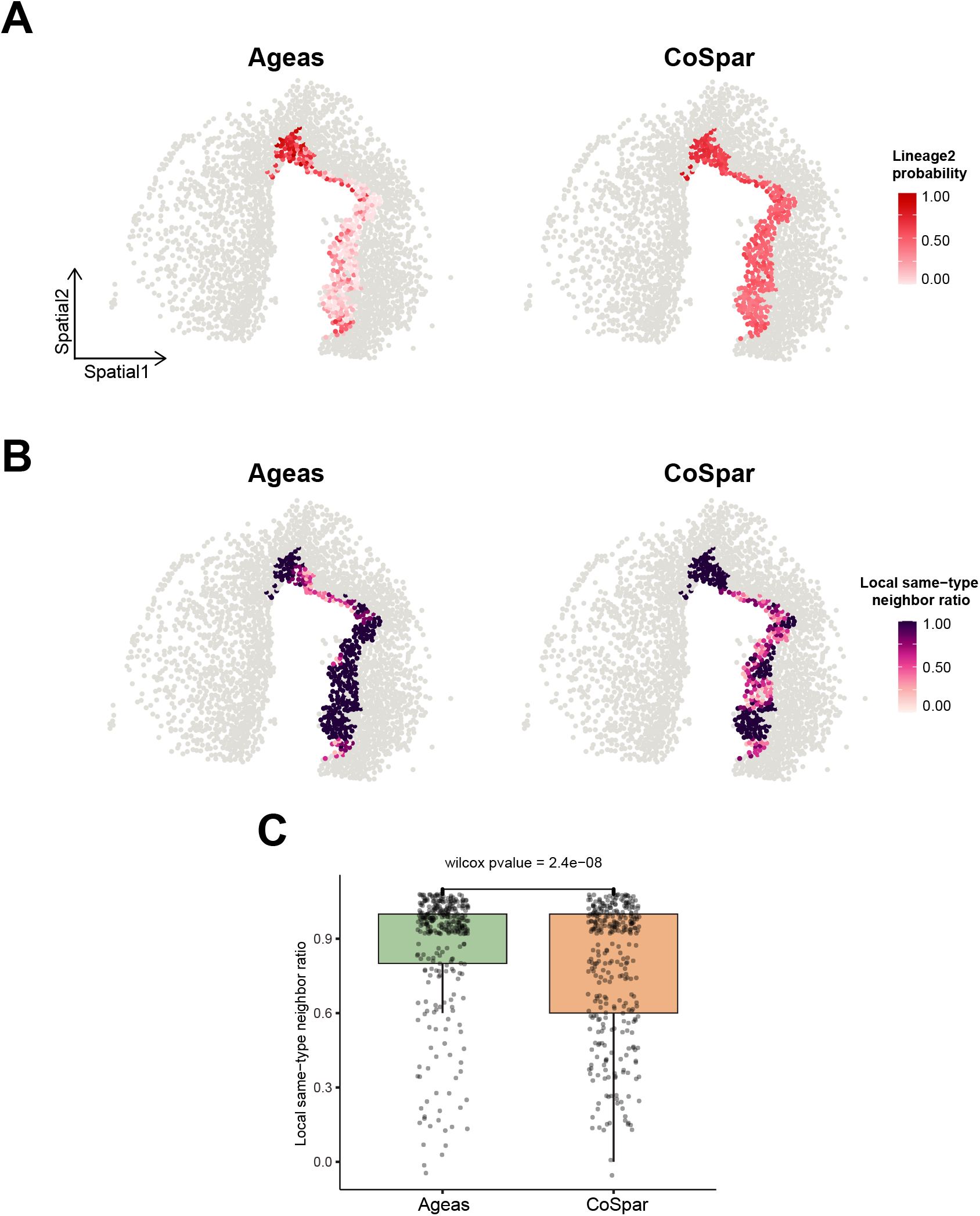
Spatial coherence quantification. **A,** Lineage2 fate probability predicted by Ageas and CoSpar. **B,** Spatial visualization of the local same-type neighbor ratio for reaEGCs based on Ageas and CoSpar predictions. Higher values indicate stronger spatial aggregation of cells sharing the same predicted fate. **C,** Box plot comparing the local same-type neighbor ratio. P-value was calculated using the two-sided Wilcoxon rank-sum test.

**Supplementary Fig. 6:**
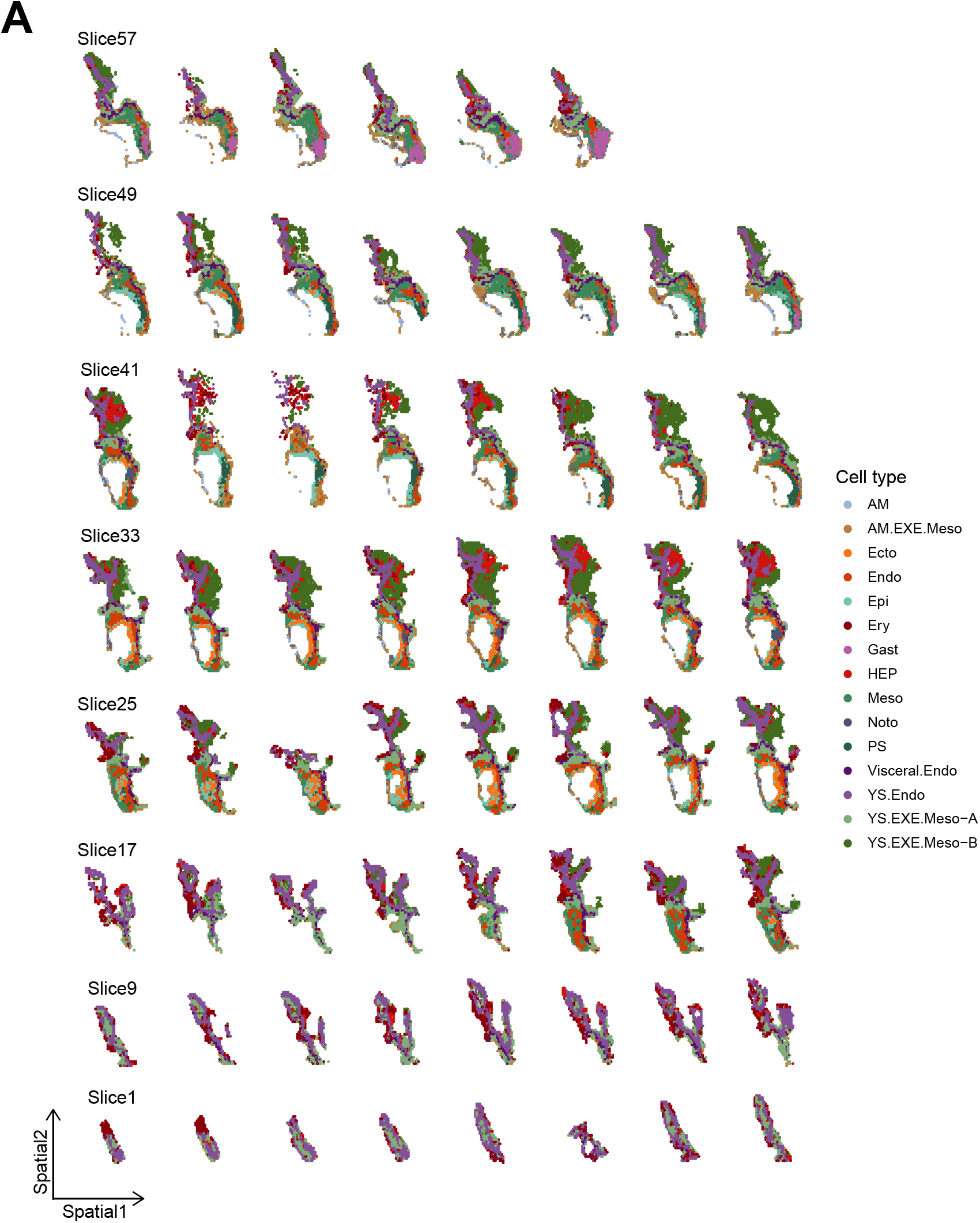
3D spatial atlas of the CS8 human embryo. **A,** Serial spatial maps showing cell type annotations for consecutive slices. The sequence proceeds from the anterior pole (Slice 1, bottom-left) to the posterior pole (Slice 64, top-right).

**Supplementary Fig. 7:**
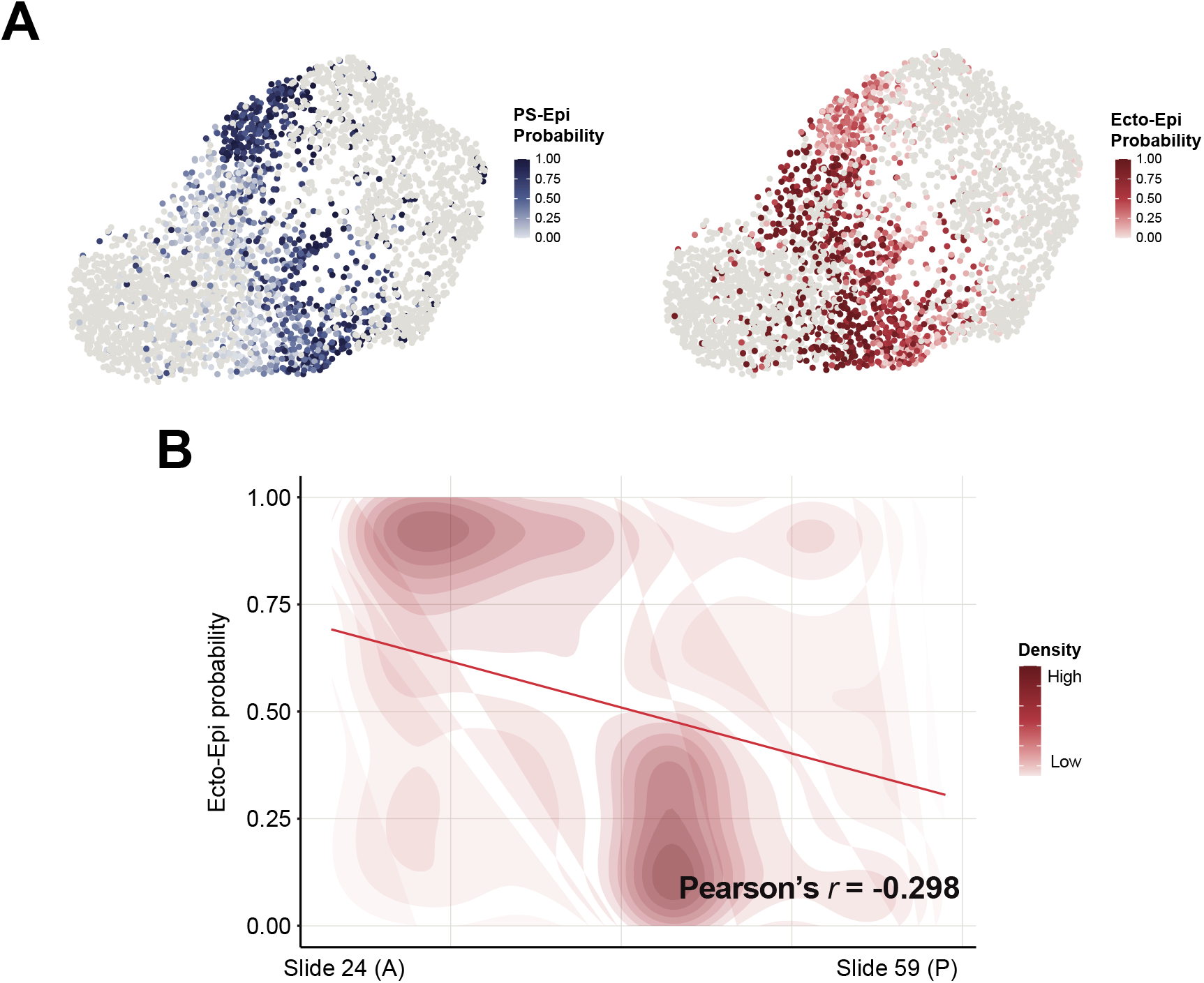
Spatial gradient of epiblast fate bias. **A,** UMAP visualization of epiblast cells colored by predicted probabilities for PS-Epi (left) and Ecto-Epi (right) fates. **B,** Correlation of predicted Ecto-Epi probabilities with the A-P spatial coordinate. Color represents the local density of epiblast cells. For instance, high density in the upper-left region indicates that anterior epiblasts predominantly exhibit high ectodermal probability.

**Supplementary Fig. 8:**
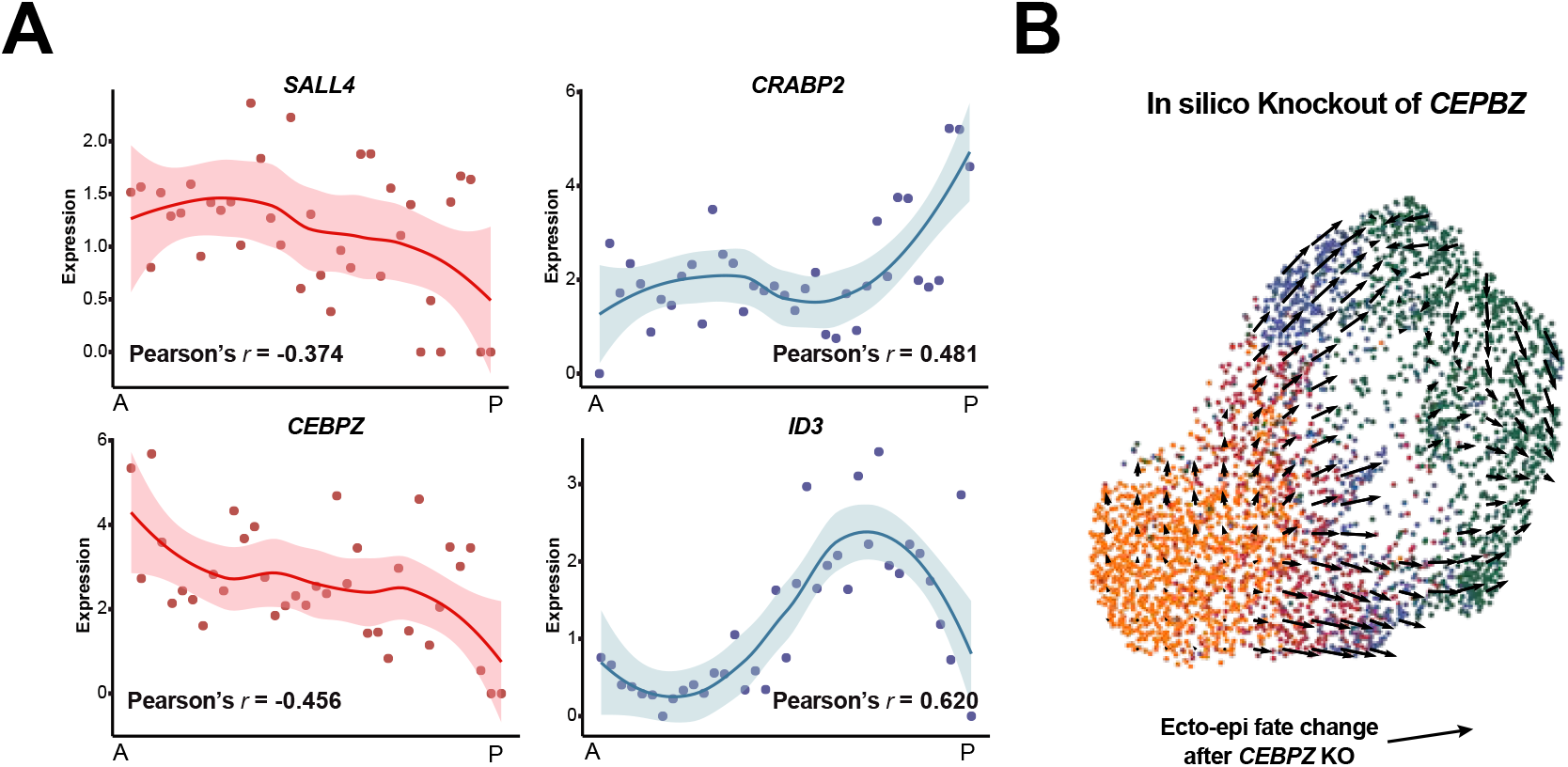
Regional molecular programs for epiblast fate specification. **A,** Spatial expression trends of representative driver genes (*SALL4*, *CRABP2*, *CEPBZ*, *ID3*) along the A-P axis. Points represent mean expression of epiblast per slice; curves show the fitted trend with Pearson correlation coefficients. **B,** CellOracle in silico perturbation analysis simulating *CEPBZ* knockout. Vector field arrows indicate the predicted shift in cell state transitions, showing a loss of ectodermal priming trajectory.

## Notes

### Competing Interest Statement

The authors have declared no competing interest.

